# Differential processing and localization of human Nocturnin controls metabolism of mRNA and nicotinamide dinucleotide metabolites

**DOI:** 10.1101/2020.01.12.903534

**Authors:** Elizabeth T. Abshire, Kelsey Hughes, Rucheng Diao, Sarah Pearce, Raymond C. Trievel, Joanna Rorbach, Peter L. Freddolino, Aaron C. Goldstrohm

**Author notes:** E.T. Abshire, University of Rochester, School of Medicine and Dentistry, Department of Biochemistry and Biophysics, 601 Elmwood Avenue, Box 712, Rochester, NY 14642, K. Hughes, Macmillan Learning, 211 E. 7th Street, Suite 300, Austin, TX 78701. These authors contributed equally (Kelsey Hughes, Rucheng Diao, Sarah Pearce). Corresponding Author: Aaron C. Goldstrohm, Mailing Address: Room 6-155 Jackson Hall, 1214A, 321 Church Street SE, Minneapolis, MN, 55455.

## Abstract

Nocturnin (NOCT) is a eukaryotic enzyme that belongs to a superfamily of exoribonucleases, endonucleases, and phosphatases. In this study, we analyze the expression, processing, localization, and cellular functions of human NOCT. We demonstrate that the NOCT protein is differentially expressed and processed in a cell and tissue type specific manner as a means to control its localization to the cytoplasm or mitochondria. Our studies also show that the N-terminus of NOCT is necessary and sufficient to confer mitochondrial localization. We then measured the impact of cytoplasmic NOCT on the transcriptome and report that it regulates the levels of hundreds of mRNAs that are enriched for components of signaling pathways, neurological functions, and regulators of osteoblast differentiation. Recent biochemical data indicate that NOCT dephosphorylates nicotinamide adenine dinucleotide (NAD) metabolites, and thus we measured the effect of NOCT on these cofactors in cells. We find that NOCT increases NAD(H) and decreases NADP(H) levels in a manner dependent on its intracellular localization. Collectively, our data indicate that NOCT can regulate levels of both mRNAs and NADP(H) cofactors in manner specified by its intracellular localization.

## Introduction

Nocturnin (NOCT) is a eukaryotic protein that belongs to the Exonuclease-Endonuclease-Phosphatase (EEP) superfamily of enzymes and is conserved from insects to vertebrates (1-6). While members of the EEP superfamily can act on diverse substrates, ranging from nucleic acids to phospholipids, NOCT is most similar to the CCR4 subclass of EEP enzymes, which is indicative of a role in RNA metabolism (1,2). CCR4 and its orthologs are hydrolytic exoribonucleases that degrade the 3′ poly-adenosine tail of mRNA substrates (i.e., they are deadenylases) (7). Indeed, sequence and structural homology of the C-terminal EEP domain of NOCT revealed that it most closely resembles CCR4 class enzymes CNOT6 and PDE12 (1,6). Nonetheless, NOCT possesses unique features that distinguish it from other CCR4 type enzymes. First, NOCT has a unique N-terminus whose function was poorly understood. Second, NOCT has a conserved, highly basic cleft adjacent to its active site that was proposed to participate in substrate recognition (1).

Phenotypic analysis of NOCT knockout mice indicates roles in cellular differentiation and metabolism. NOCT was initially discovered as a transcript whose expression is controlled by circadian rhythm in the retina of *Xenopus* (2). Subsequent analysis in mice revealed that while NOCT expression is in part influenced by circadian rhythms, NOCT is not required for circadian gene expression or behavior (4). Instead, knockout of NOCT results in resistance to high fat diet induced obesity. NOCT knockout mice are viable but have defects in absorption, transport, and storage of fat (8). These mice exhibit resistance to hepatic steatosis and have greatly diminished adipose tissue (4). In addition, NOCT knockout mice have increased bone mass with reduced bone marrow adiposity, indicating that NOCT negatively regulates osteogenesis while promoting adipogenesis, which was further supported by cell-based assays (4,9,10). The biological roles of NOCT in humans remain largely unknown, and the molecular functions of NOCT remain elusive. Addressing this challenge is essential in order to understand how NOCT controls metabolism and cellular differentiation.

Given its relationship to CCR4-type deadenylases, substantial effort has focused on assessing the ability of NOCT to degrade RNA substrates and its potential impact on gene expression. Initial biochemical assays suggested that NOCT could degrade poly(A) RNA in vitro; however, subsequent analyses using pure recombinant NOCT did not detect hydrolysis of poly(A) substrate RNAs (1,2,11,12). Thus, while NOCT does not degrade poly(A) RNA on its own, it remains plausible that NOCT is an exoribonuclease that may require obligatory, unknown partners, cofactors, or modifications. Additionally, because only a few RNA substrates have been tested, it is possible that NOCT may have specific RNA sequence and/or structure requirements.

Multiple approaches support the model that NOCT can act on mRNAs in vivo. First, when directed to a reporter mRNA in cell-based, tethered function assays, NOCT causes translational inhibition and degradation of that mRNA. The repressive activity of NOCT is dependent on the 3′ end of the mRNA; it can affect poly(A) reporter mRNA but not a derivative with a 3′ triple helix structure. Moreover, mutations in conserved residues of the putative active site of NOCT reduced its repressive activity (1). In these aspects, NOCT exhibits mRNA regulatory activity similar to the CCR4-NOT deadenylase complex (7). Biochemical analysis of NOCT-protein interactions provided further support for a role in mRNA metabolism. Characterization of *Drosophila* and mammalian NOCT protein complexes indicated that they associate with components of the CCR4-NOT deadenylase complex, which initiates the major mRNA decay pathway (13,14). Further evidence in support of NOCT-mediated mRNA regulation comes from studies that employed NOCT knockout mice or RNAi-mediated depletion of NOCT in mouse cells; loss of NOCT expression resulted in changes in gene expression at the transcript level, and therefore may reflect action of NOCT on mRNA substrates (15-18). In one instance, NOCT was reported to physically associate with a target mRNA (i.e. specific isoforms of Igf1 mRNA), supporting the hypothesis that NOCT has a direct effect on that mRNA (10). The effect of human NOCT on mRNAs remained unexplored; therefore, in this study we analyzed its impact on the transcriptome in cultured human cells.

Homology of NOCT to EEP enzymes suggests that NOCT may act on other types of phosphorylated substrates instead of, or in addition to, mRNA. Precedent for this hypothesis is demonstrated by the CCR4-type deadenylase PDE12, which hydrolyzes the signaling molecule 2′-5′ oligo-adenylate, in addition to poly(A) RNA substrates with 3′-5′ phosphodiester bonds (19-21). In our previous work, we analyzed the ability of NOCT to act on a library of phosphorylated nucleotides, sugars, lipids, none of which were hydrolyzed by pure recombinant NOCT (1). Interestingly, a recent study reported that purified NOCT can remove the 2′ phosphate of the nicotinamide adenine dinucleotide phosphate cofactors NADPH and NADP^+^ (22). In this study, we explore the potential effect of human NOCT on the levels of nicotinamide adenine dinucleotide cofactors and ATP in cells.

The intracellular localization of NOCT remains incompletely understood and could have a profound effect on its molecular function and substrates. Previously, over-expressed, epitope tagged NOCT was reported to be either cytoplasmic or peri-nuclear, whereas endogenous NOCT has been reported to localize to the nucleus or cytoplasm of cells in mouse embryos or to the cytoplasm in *Xenopus* retina (2,9,22,23). Intriguingly, sequence analysis of the unique N-terminus of mouse and human NOCT detected the presence of a putative mitochondrial targeting sequence (**Figure 1a**) (6). Moreover, we observed that the N-terminus of NOCT possesses two potential translation initiation sites, one of which would include the mitochondrial targeting sequence and one which would exclude it. Therefore, we examined the intracellular localization of NOCT and potential for differential translation initiation to produce protein isoforms. We also explore the tissue specific expression pattern of human NOCT protein.

**Figure 1.**
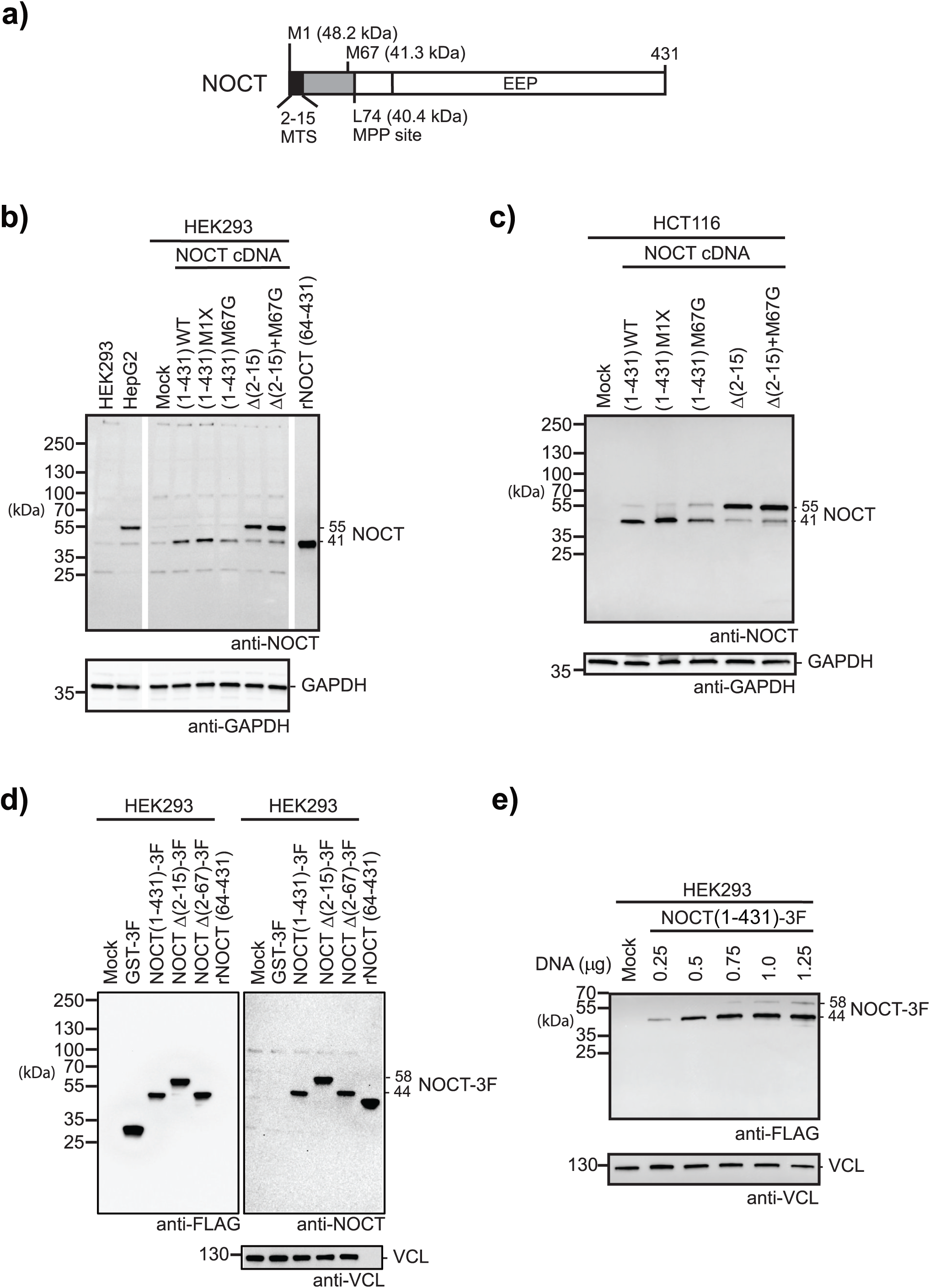
NOCT protein is post-translationally processed in a manner consistent with mitochondrial targeting. **a)** Diagram of the NOCT protein coding sequence. The locations of a predicted Mitochondrial Targeting Sequence (MTS) and Exo-Endo-Phosphatase (EEP) domain are indicated. Two potential translation sites are indicated at the top (methionine codons M1 and M67). The location of a predicted Mitochondrial Processing Peptidase (MPP) cleavage site, L74, is indicated at the bottom. Predicted molecular weights of the resulting NOCT isoforms are indicated. **b)** Western blot analysis of translation initiation of NOCT protein. Endogenous NOCT protein was detected in cell extracts from human embryonic kidney (HEK293) or human liver (HepG2) cell lines, indicated at the top. The apparent molecular weights of endogenous NOCT protein were observed at 41 kDa in HEK293 cells and 55 kDa in HepG2 cells. NOCT protein expression in HEK293 cells was also analyzed in cells transiently transfected with wild type (WT) NOCT cDNA encoding amino acids 1-431. The role of the two potential initiation sites was tested by mutating M1 AUG codon to GGG (M1X) or M67 AUG codon to a glycine codon, GGG (M67G). The role of the N-terminal predicted MTS in processing of NOCT protein was tested; deletion of the MTS in NOCT Δ(2-15) or NOCT Δ(2-15)+M67G abolishes processing, consistent with mitochondrial localization and processing. “Mock” indicates untransfected cells. For all western blots, equal mass of protein from cell lysates was loaded in each lane of the gel. NOCT protein was detected using western blotting with antigen affinity purified, rabbit polyclonal α-NOCT antibody. Recombinant purified NOCT, rNOCT (64-431), served as a positive control for western blot detection. Blots were probed using α-GAPDH antibody to assess equivalent loading of the samples. Molecular weight markers in kilodaltons (kDa) are indicated on the left. Apparent molecular weights are indicated on the right. **c)** The same approach as in b) except using human colon carcinoma cell line, HCT116. **d)** Processing of NOCT protein requires the MTS. Expression of NOCT in HEK293 cells from transfected plasmid that expresses the coding sequence fused to C-terminal 3xFLAG epitope tag. NOCT (1-431)-3F is processed to produce a 44 kDa product, including 3 kDa of additional mass from the C-terminal 3xFLAG tag. Deletion of the MTS in NOCT Δ(2-15)-3F results in expression of a 58 kDa form of NOCT-3F, and deletion of amino acids 2-67 resulted in expression of NOCT Δ(2-67)-3F of 44 kDa. Equal mass of protein from cell lysates was analyzed for each sample. NOCT was detected using western blotting with α-FLAG (left panel) and rabbit α-NOCT (right panel). Western blot of vinculin using α-VCL assess equivalent loading of the lanes. Molecular weight markers in kilodaltons (kDa) are indicated on the left. Apparent molecular weights are indicated on the right. **e)** Detection of processed (44 kDa) and unprocessed (58 kDa) NOCT in HEK293 cells transfected with titration of plasmid expressing full length NOCT (1-431)-3F in HEK293 cells. The amount of NOCT expression vector is indicated at the top. NOCT was detected using western blotting with rabbit α-FLAG. Equivalent mass of protein from each cell lysate was analyzed in each lane of the gel. Blots were reprobed using α-VCL to assess equivalent loading of the gel lanes.

In this report, we demonstrate that NOCT protein is differentially processed in a cell type specific manner to control its localization to the cytoplasm and mitochondria. The N-terminus of NOCT is necessary and sufficient to confer mitochondrial localization. Analysis of human and mouse tissues provides evidence for tissue type specific control of processing of NOCT protein. By employing a gain of function approach with transcriptome-wide quantitation, we then demonstrate that the cytoplasmic form of NOCT regulates the levels of hundreds of mRNAs that are enriched for components of signaling pathways and neurological functions. Finally, we explore the impact of NOCT on nicotinamide adenine dinucleotide metabolites and observe specific effects on NADPH and NADP^+^. Together, our data indicate that NOCT can control levels of both mRNAs and NADP(H) in manner specified by its intracellular localization.

## Results

### Nocturnin protein is post-translationally processed in a tissue specific manner

To analyze NOCT protein expression, we generated a polyclonal antibody that was then antigen affinity-purified and validated by western blot detection of over-expressed NOCT (encoding amino acids 1-431) in HEK293 cells and recombinant purified human NOCT protein (64-431) (**Figure 1b**). Western blotting of equal amounts of cell extracts from HEK293 and HepG2 cell lines resulted in the detection of two NOCT protein isoforms (**Figure 1b**). The major anti-NOCT reactive band detected in the HepG2 liver cell line migrates with an apparent molecular weight of ∼55 kDa, which is slightly larger than the predicted 48.2 kDa for full length 431 amino acid NOCT protein (**Figure 1a and 1b**). We note that the N-terminus of NOCT is highly basic (see below), which may contribute to the difference in electrophoretic mobility. This 55 kDa band was not observed in HEK293 cell extract. In both HEK293, and HepG2 cells, a ∼41 kDa protein was also detected, albeit with low relative abundance. The observation of two NOCT protein isoforms suggested the potential for differential translation initiation, protein processing, or alternative mRNA processing. Alternative NOCT mRNA isoforms have not been reported in humans and we have not detect other isoforms in human RNA-seq data; therefore, we focused on the potential for differential translation initiation and/or protein processing.

Examination of the NOCT mRNA suggested the potential for two translation initiation sites: M1 and M67 (**Figure 1a**). Translation typically initiates at the first AUG codon (M1) with the optimal Kozak consensus sequence context (5′-**GCCRCC<u>AUG</u>G**) that is encountered by 43S ribosomal pre-initiation complexes during scanning of the mRNA (24-26). For NOCT, initiation at M1 would produce the full length 431 amino acid protein. However, the sequence context of M1 (5′-C**CCG**G**C<u>AUG</u>**U, bold indicates match to the Kozak consensus) differs from the optimal Kozak sequence, raising the possibility for leaky scanning that might lead to initiation at the downstream M67 (5′-UGUU**CC<u>AUG</u>G**). To interrogate utilization of these initiation sites, we cloned the NOCT cDNA, including its natural 5′ leader sequence and the open reading frame encompassing amino acids 1-431, into an expression vector. We then mutated either initiation codon from AUG to GGG (M1X or M67G) and transfected the constructs into HEK293 cells. Western blot analysis indicates that M1 is the predominant initiation site (**Figure 1b**), whereas M67 can be utilized in the construct wherein M1 is lost (**Figure 1b**). Expression of wild type NOCT cDNA (1-431) increased the level of the ∼41 kDa form in HEK293 cells. Mutation of M1 produced a protein of similar apparent size, consistent with the scanning mode of translation wherein M67 now becomes the first AUG codon and therefore serves as the site of translation initiation in the absence of M1. We also mutated M67 alone and observed that this construct produced a ∼41 kDa product. These observations support the hypothesis that translation of wild type NOCT protein initiates at M1, resulting in expression of a ∼55 kDa pre-protein that is then processed into the 41 kDa form in HEK293 cells.

Analysis of the NOCT amino acid sequence detected a putative N-terminal mitochondrial targeting presequence and cleavage site for the mitochondrial processing peptidase (MPP) (**Figure 1a**), which are conserved in several vertebrates (6,27). The MitoFates and TPpred2 algorithms score the probability that the NOCT N-terminus has a mitochondrial presequence at 0.94 and 0.98, respectively (27-29). Mitochondrial targeting signals typically reside in the first 90 N-terminal residues, form amphipathic helices, have a high arginine content, and contain few negatively charged residues (27). Consistent with these features, amino acids 2-15 are predicted to be an amphipathic alpha-helix and the first 74 amino acids have an isoelectric point of 12.1, with 12 arginines and only one negatively charged residue (6). Deletion of amino acids 2-15 reduced the MitoFates probability score from 0.94 to 0.035. MitoFates additionally predicts a consensus cleavage site for MPP, including the Arg at −2 position relative to the cleavage site after L74 (**Figure 1a**) (27). The resulting NOCT isoform (aa 75-431) would have a molecular weight of 40.4 kDa, consistent with the observed ∼41 kDa band. Based on these features, the NOCT pre-protein is predicted to be recognized by the translocase of outer membrane (TOM), facilitating import of NOCT pre-protein into the mitochondrial matrix where it is then cleaved by MPP to produce the ∼41 kDa isoform (30). As predicted, deletion of the MTS sequence, NOCT Δ(2-15), abrogated processing of NOCT and resulted in accumulation of a ∼55 kDa protein in HEK293 cells (**Figure 1b**). The M67G mutation did not alter expression and processing of this form of NOCT. Together, these results are consistent with initiation at M1 followed by MTS dependent processing of NOCT from the ∼55 kDa form to the ∼41 kDa form. We corroborated these observations in the HCT116 colon cancer cell line (**Figure 1c**), supporting the utilization of M1 during translation initiation to produce a polypeptide containing an MTS that facilitates proteolytic processing of NOCT. Interestingly, the processing appears to be cell-type specific, as HepG2 liver cells predominantly produce an unprocessed ∼55 kDa form of NOCT (**Figure 1b**).

To further corroborate the processing of NOCT protein, we appended a 3xFLAG (3F) tag to the C-terminus of NOCT and examined the proteins produced in HEK293 cells from plasmids that express wild type NOCT (1-431) or the deletion of MTS (NOCT Δ(2-15)) or N-terminus (NOCT Δ(2-67)). Full length NOCT (1-431)-3F was processed to form a ∼44 kDa product, whereas deletion of the MTS in NOCT Δ(2-15)-3F prevented processing, resulting in expression of a ∼58 kDa product (**Figure 1d**). A larger deletion, NOCT Δ(2-67)-3F which lacks the N-terminal sequence preceding and M67, produced the expected ∼44 kDa product. These constructs are therefore processed to form products with apparent molecular weights consistent with the sizes for these NOCT constructs with an additional 3 kDa of apparent molecular weight from the 3x FLAG epitope tag. Therefore, we observe efficient processing of NOCT in HEK293 cells using western blotting with either anti-FLAG or anti-NOCT antibodies, consistent with mitochondrial import and subsequent proteolytic cleavage. We also titrated NOCT expression plasmid and observed that at higher amounts of transfected plasmid, the precursor ∼58 kDa form becomes evident, along with the corresponding increase in processed ∼44 kDa NOCT (**Figure 1e**).

We then investigated the role of MTS-dependent processing of NOCT in localization to mitochondria. We transfected the NOCT expression constructs into 143B human osteosarcoma cells, which are well-suited to microscopy studies, and visualized the localization of each FLAG-tagged construct using immunofluorescence (**Figure 2a**). The cells were also stained with MitoTracker and DAPI dyes to visualize mitochondria and nuclei, respectively. As a negative control, we transfected a FLAG-tagged glutathione S-transferase (GST-3F) expression construct, which was localized throughout the cytoplasm. The NOCT (1-431)-3F construct colocalized with the signal from the MitoTracker stain, indicating that the 143B cell line localizes NOCT to the mitochondria. To determine if mitochondrial localization is dependent on the MTS, we tested the NOCT Δ(2-15)-3F construct, which is resistant to processing (**Figure 1**) and prevented the mitochondrial localization of NOCT, as the staining of this construct was diffuse and cytoplasmic (**Figure 2a**). We also visualized the localization of NOCT Δ(2-67)-3F, which also was expressed throughout the cytoplasm. These data indicate that wild type NOCT is localized to the mitochondria, dependent on the N-terminal MTS and consistent with the NOCT protein processing data shown in **Figure 1**.

**Figure 2.**
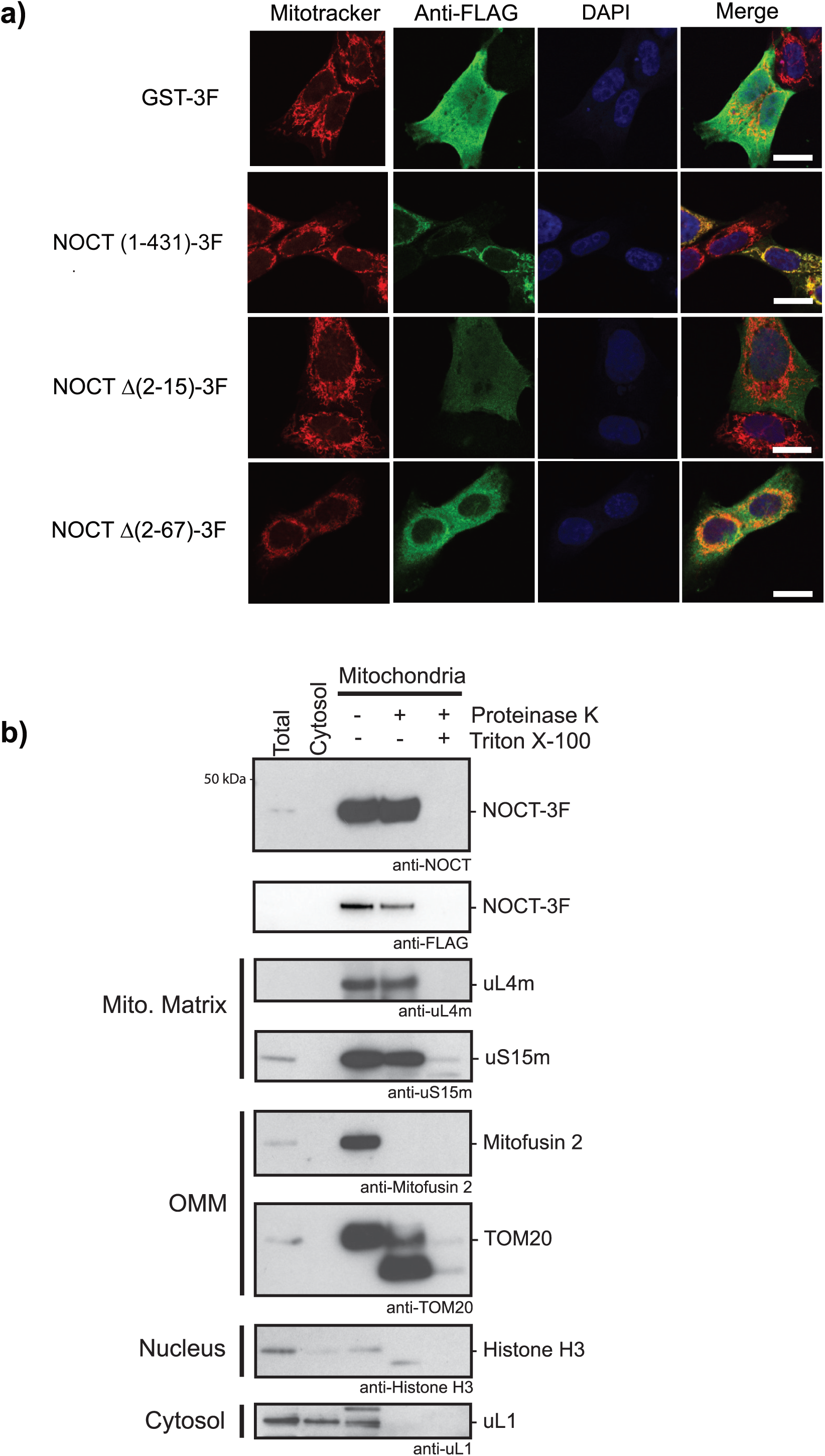
NOCT protein is localized to the mitochondria, dependent on its N-terminal mitochondrial targeting sequence. **a)** Immunofluorescence analysis of intracellular localization of NOCT constructs. NOCT-3F is localized to the mitochondria whereas NOCT Δ(2-15)-3F and NOCT Δ(2-67)-3F constructs, which lack the MTS, are predominantly localized to the cytoplasm. The NOCT expression plasmids indicated on the left were transfected into 143B human osteosarcoma cells and visualized using anti-FLAG monoclonal antibody immunofluorescence against the C-terminal 3x-FLAG epitope on each construct with Alexa Fluor Plus 488 secondary antibody (green). Mitochondria are visualized with MitoTracker Red CMXRos (red) and nuclei are visualized with DAPI (blue). GST-3F served as a control. The white scale bar is 10 µm. **b)** NOCT fractionates with the mitochondria, as tested by western blotting of subcellular fractions (including total, cytosol, and mitochondria) prepared from HEK293T cells that were transfected with NOCT (1-431)-3F. The mitochondrial fractions were divided into three equal aliquots and were either left untreated or were treated with Proteinase K or Proteinase K with Triton X-100. The fractions were then analyzed by SDS PAGE and western blotting with the indicated antibodies. NOCT was detected using guinea pig polyclonal anti-NOCT and goat monoclonal anti-DDDDK (FLAG) antibodies. Fractionation was validated using the following antibodies: anti-uL4m and anti-uS15m for the mitochondrial matrix, anti-Mitofusin 2 and anti-TOM20 for the outer mitochondrial membrane, anti-Histone H3 for the nucleus, and anti-uL1 for the cytoplasm.

We next examined the localization of over-expressed wild type NOCT (1-431)-3F in HEK293T cells using subcellular fractionation (**Figure 2b**). Cell homogenates were separated into cytoplasmic and mitochondrial fractions and NOCT-3F was detected by western blotting with anti-NOCT or anti-FLAG antibodies. We observed that NOCT-3F was only detectable in the mitochondrial fraction with an apparent size of ∼41 kDa, consistent with processing of NOCT in an MTS-dependent manner. As expected, control cytosolic uL1 ribosome protein and nuclear histone H3 proteins were not enriched in the mitochondrial fraction. The mitochondrial fractions were subsequently treated with Proteinase K to remove contaminating proteins outside of intact mitochondria. Under these conditions, outer mitochondrial membrane proteins Mitofusin 2 and TOM20 undergo proteolysis due to their exposure to Proteinase K, whereas mitochondrial matrix proteins uL4m and uS15m, both constituent proteins of the mitochondrial ribosome, are protected from proteinase activity. The ∼41 kDa NOCT was also protected from proteolysis, indicating that it is protected within the mitochondrial membranes. In contrast, when the membranes were disrupted by treatment with Triton X-100, proteolysis of NOCT occurred, in a manner similar to matrix proteins uL4m and uS15m. Together, our data indicate that NOCT can be localized to the mitochondria in HEK293T and 143B cell types, where it is imported and processed into a ∼41 kDa form.

To ascertain if the N-terminus of NOCT is sufficient to confer proteolytic processing and mitochondrial localization, we fused the first 86 N-terminal amino acids of NOCT to enhanced green fluorescent protein (EGFP) and expressed the construct in HEK293 cells. A minor band was observed with an apparent molecular weight of ∼42 kDa, which is higher than the predicted 35.7 kDa of the NOCT(1-86) EGFP pre-protein and, like full length NOCT, is likely the result of the highly basic nature of the N-terminus (**Figure 3a**). A major band with apparent molecular weight of ∼26 kDa was also produced, consistent with MPP processing of the NOCT mitochondrial presequence (predicted molecular weight 27.9 kDa), and slightly larger than the ∼25 kDa EGFP protein, which consistent with the additional 12 amino acids remaining after MPP-mediated cleavage. To determine if the NOCT N-terminus EGFP fusion protein can localize to the mitochondria, we visualized the localization of this protein using fluorescence microscopy. When expressed in human osteosarcoma cells, the NOCT (1-86) EGFP protein colocalized completely with the MitoTracker stain and was not observed throughout the cell body or nucleus (**Figure 3b**). Together, our data indicate that the NOCT N-terminal 86 amino acids are sufficient for processing and localization of EGFP fusion protein to the mitochondria.

**Figure 3.**
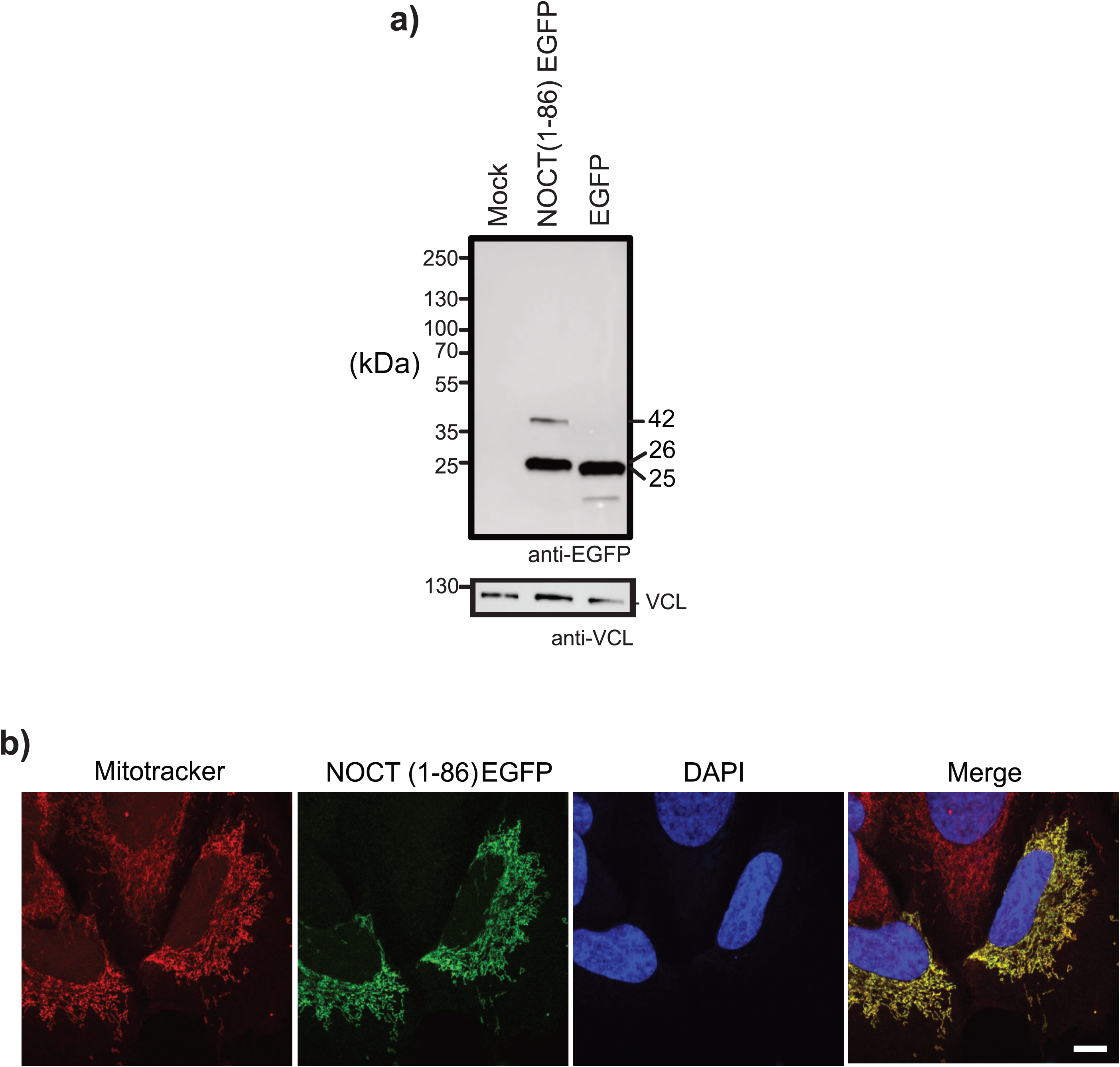
The N-terminus of NOCT is sufficient for processing and mitochondrial localization. **a)** The N-terminus of NOCT is sufficient to confer protein processing consistent with mitochondrial localization. The N-terminal amino acids 1-86 of NOCT were fused to the N-terminus of EGFP to create NOCT (1-86) EGFP expression construct, which was then transfected into HEK293 cells. Cells transfected with EGFP construct served as a control. Equal amounts of cell lysates were then analyzed by SDS PAGE and western blotting using anti-EGFP antibody or, as a loading control, anti-Vinculin (VCL). Molecular weight markers in kilodaltons (kDa) are indicated on the left. Apparent molecular weights are indicated on the right. **b)** NOCT (1-86) EGFP protein (green) is localized to the mitochondria in 143B cells. Mitochondria are visualized with MitoTracker (red) and nuclei are visualized with DAPI (blue). The white scale bar is 10 µm.

We observed that NOCT protein appears to be differentially processed in HEK293 relative to NOCT expressed in HepG2 cells (**Figure 1b**), suggesting that processing of NOCT may be cell type specific. While the expression pattern of NOCT in human tissues has not been investigated in detail, available RNA-seq data indicates broad expression of NOCT mRNA in human tissues (**Supporting Information Figure S1**). Human NOCT mRNA is most highly expressed in adipose, breast, liver, lung, muscle, kidney, and certain regions of the brain (e.g. cerebellum, frontal cortex), whereas the lowest NOCT expression occurs in ovary, pancreas, bladder and other brain regions (e.g. spinal cord, amygdala, basal ganglia).

Using our NOCT antibody, we examined endogenous NOCT protein expression in a collection of human tissues in order to observe the extent to which processing of NOCT appears to be tissue-specific. The NOCT protein, with apparent molecular weight of ∼55 kDa, was detected in human brain, kidney, and liver, indicating that full length, unprocessed NOCT is expressed in multiple tissue types. The observation of ∼55 kDa NOCT in liver is consistent with the form we observed in HepG2 liver cell line in **Figure 1b**. Several other NOCT species were also observed, including a band at ∼40 kDa in brain, skeletal muscle, and to a lesser degree in heart, suggesting that those tissues can process the NOCT pre-protein in a manner consistent with mitochondrial localization and MPP-mediated processing. We note that bands were detected at ∼80 kDa band in brain and a ∼16 kDa band in small intestine; however, the origins of these bands cannot currently be accounted for and may result from cross-reaction of the anti-NOCT antibody. NOCT protein expression was below the limit of detection in lung, stomach, spleen, ovary and testes.

We also investigated NOCT protein expression patterns in mouse tissues (**Figure 4b**). The RefSeq annotated mRNA encoding full-length murine NOCT protein (aa 1-429) is expected to produce a protein with predicted molecular weight of 48 kDa with an N-terminal MTS (aa 2-15) and MPP cleavage site (L72) embedded in a presequence with 11 arginines and a pI of 12.2 (MitoFates P = 0.73). We therefore expect to observe that murine NOCT is processed similarly to the human enzyme. Indeed, a form of NOCT with apparent molecular weight of ∼55 kDa is observed in brain, lung, and stomach, while a major band with apparent molecular weight of ∼40 kDa was detected in brain, heart, and skeletal muscle, and may correspond to MPP-processed NOCT. An additional ∼48 kDa band is also detected in brain, heart, small intestine, kidney, liver, spleen, testis (**Figure 4b**). We note that a second mouse NOCT mRNA isoform (XM_006500956.1) is annotated in the RefSeq database and encodes a 372 aa protein (42.2 kDa)(XP_006501019). Transcription of this alternative mRNA isoform initiates within the intron 1 of the NOCT gene and produces an mRNA with a unique 5′ exon that encodes an alternative 4 amino acids (MALP) in place of the first 61 amino acids normally encoded within exon 1. The ∼48 kDa band therefore may represent NOCT translated from this mRNA transcript (**Figure 4b**). Additional anti-NOCT reactive ∼66, ∼80, and ∼97 kDa bands are detected in brain and small intestine, which cannot be accounted for by documented transcripts. In addition to processing, it is also possible that post-translational modifications contribute to the apparent sizes of the different observed NOCT species in tissues where NOCT is detected, though no such modifications have been specifically described. In summary, we have demonstrated that the N-terminus of NOCT contains an MTS that is both necessary and sufficient for processing of NOCT, and that differential expression and processing of endogenous NOCT occurs in a tissue-dependent manner.

**Figure 4.**
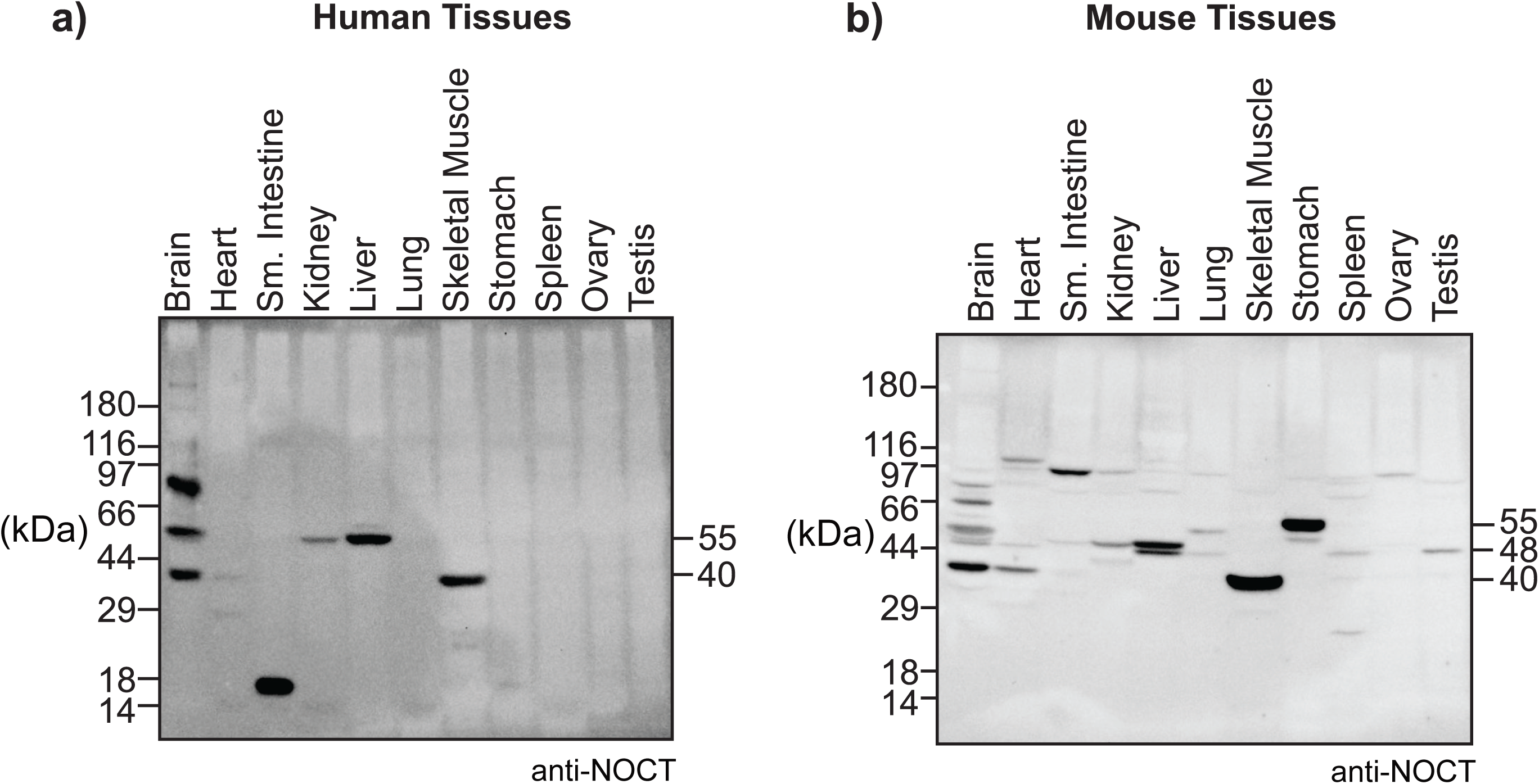
NOCT protein is differentially expressed in human and mouse tissues. Western blotting of human **a)** and mouse **b)** tissues, indicated at the top, was performed using rabbit polyclonal anti-NOCT antibody. Commercially produced tissue blots contain 50 µg of each tissue. Bands in the 40-55 kDa range are considered feasible to be NOCT. Molecular weight markers are indicated on the left. Apparent molecular weights of NOCT bands detected in human brain, heart, kidney, liver, and skeletal muscle, and in mouse brain, heart, kidney, liver, lung, skeletal muscle, stomach, spleen, and testis are indicated on the right.

### Impact of cytoplasmic NOCT on the transcriptome

We next investigated the effect of the cytoplasmic localized human NOCT on transcriptome-wide RNA levels. To do so, we generated HEK293 cell lines that stably over-express NOCT Δ(2-15)-3F construct, which lacks the MTS and is localized to the cytoplasm. We note that the human embryonic kidney cell line, HEK293, has characteristics of kidney, adrenal, and neuronal tissues, and its use was advantageous because it expresses low levels of endogenous, processed ∼41 kDa NOCT (**Figure 1b**) (31,32). As a negative control, we also generated cells that stably express glutathione S-transferase-3x FLAG (GST-3F). Three independent clonal cell lines were generated for each construct and served as biological replicates (**Figure 5a**). It is noteworthy that we also attempted to analyze loss of function of the NOCT; however, we were unable to obtain viable, homozygous knockout cells generated by CRISPR/Cas9 mediated approaches, despite repeated attempts in multiple human cell lines.

**Figure 5.**
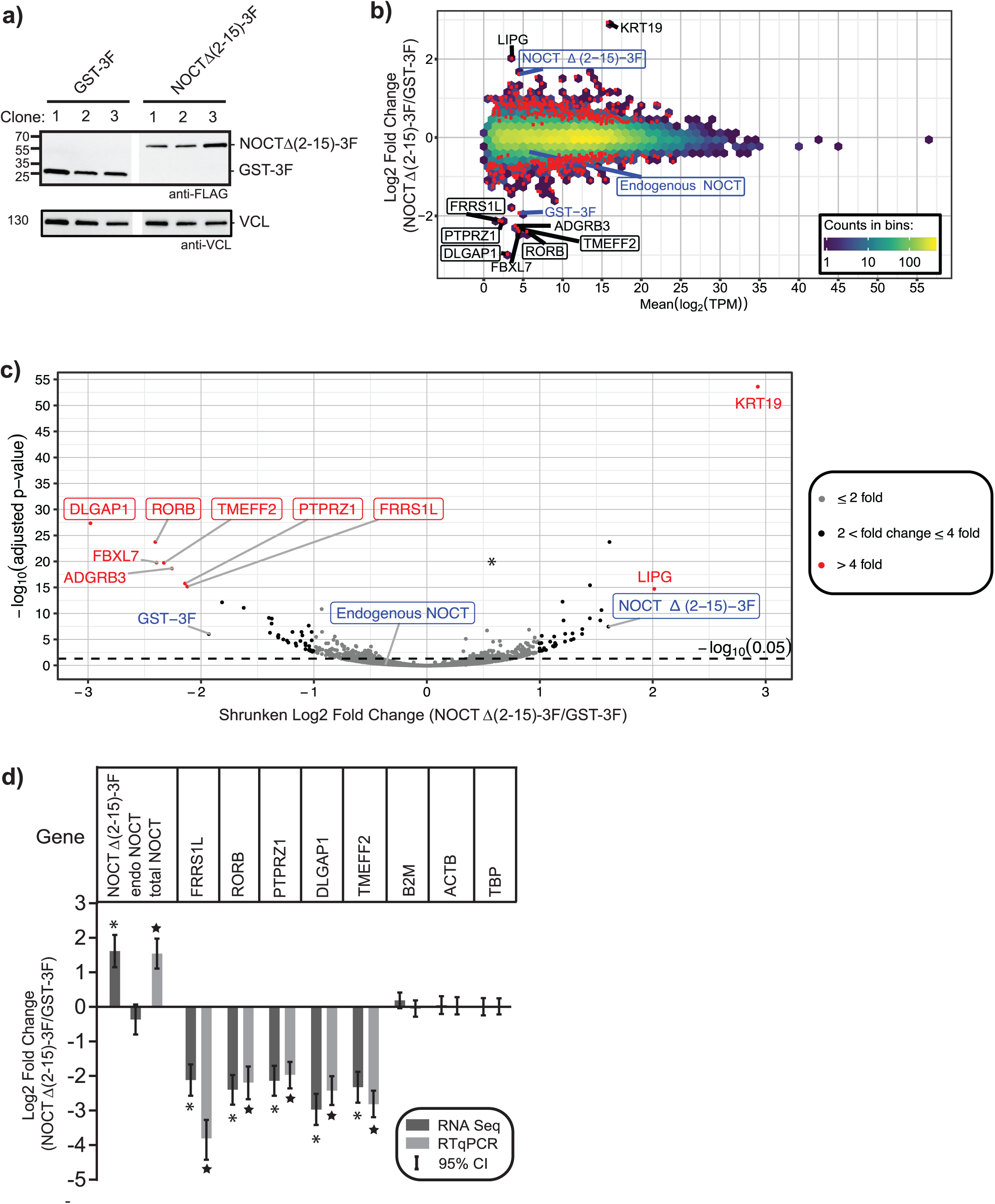
Impact of cytoplasmic NOCT Δ(2-15)-3F on the transcriptome. **a)** GST-3F or NOCT Δ(2-15)-3F proteins were stably transfected and expressed in three separate clonal HEK293 cell lines, as assessed by western blotting of equal mass of each cell extract using α-FLAG. Western blot detection of vinculin (VCL) confirmed equivalent loading of the gel lanes. **b)** Differential gene expression was measured by performing RNA-seq on rRNA-depleted RNA isolated from the three clonal HEK293 cell lines that over-expressed NOCT Δ(2-15)-3F relative to the three negative control HEK293 cell lines that expressed GST-3F. Gene expression changes were analyzed using DESeq2. Log2 fold changes of gene expression were plotted relative to the log2 average relative expression level of all six samples, measured in transcripts per million reads (TPM). The scaled color shows the counts of genes in each hex-bin. Red points indicate genes with significant changes in gene expression by an adjusted p-value threshold of 0.05. The black labels indicate genes with expression changes greater than or equal to four-fold. In panels b and c, blue labels point to the values for over-expressed NOCT Δ(2-15)-3F and the endogenous NOCT and the labelled boxes indicate genes for which we obtained qPCR validation. **c)** Volcano plot comparing statistical significance of measurements versus the log2 fold change in gene expression between NOCT Δ(2-15)-3F and GST-3F conditions. The red points and labels show the genes that have a greater than or equal to four-fold change, while black points are for genes with less than four-fold changes but at least two-fold changes, and grey points for genes with less than two-fold changes. The dashed line is the FDR-corrected p-value threshold of 0.05. All data and statistics for the RNA-seq analysis are reported in Supporting Information Table S1. **d)** Corroboration of NOCT-mediated regulation of FRRS1L, RORB, PTPRZ1, DLGAP1, and TMEFF2 mRNAs by RT-qPCR. The expression level of each gene was measured in RNA isolated from 3 replicate samples from each of the 3 clonal HEK293 cell lines expressing either NOCT Δ(2-15)-3F or GST-3F. The log2 fold change of each gene was then calculated as described in the Experimental Procedures and plotted along with the log2 fold changes measured in the RNA-seq analysis. The 95% credible intervals for each measurement are indicated above and below the mean log2 fold change values. The “*” symbol indicates that the observed fold change measured in the RNA-seq assay has FDR-corrected p value of <1×10-8. For the RT-qPCR measurements, the “★” symbol indicates that there is a >95% posterior probability of a change of two-fold. The B2M, ACTB, and TBP mRNAs were also measured and where unchanged in both RNA-seq and RT-qPCR assays. Data and statistics are reported in Supporting Information Table S2.

To analyze changes in RNA abundances induced by NOCT, total RNA was isolated from each cell line, ribosomal RNA was depleted, and then stranded, whole transcriptome RNA sequencing was performed. The resulting data measured 21,486 features in the three replicates for each of the different cell lines, including 21,419 Ensembl genes and 67 custom features like over-expression ORFs and ERCC spike-ins. Using stringent statistical significance thresholds (adjusted p-value ≤ 0.05) for expression changes, we identified 490 genes that are significantly affected by NOCT over-expression, with log2 fold changes ranging from −2.97 to 2.94 (**Figure 5b**). The affected genes spanned a broad range of expression levels, though genes affected to the largest degree were within the lower end of the expression range (**Figure 5b**). Consistent with the experimental design, NOCT Δ(2-15)-3F was over-expressed by 3.1-fold. NOCT Δ(2-15)-3F caused a decrease in expression of 235 genes relative to the control cells, including 31 genes that were negatively regulated between 2 and 4-fold and 7 genes downregulated by greater than 4-fold (**Figure 5c**). Another 255 were upregulated in cells expressing NOCT Δ(2-15)-3F, with increased levels of 32 genes between 2 and 4-fold and 2 genes with greater than 4-fold changes. **Supporting Information Table S1** provides a complete list of all the measured and affected transcripts, along with statistical parameters.

We corroborated the RNA-seq measurements by measuring the changes in mRNA expression levels of select mRNAs using reverse transcription coupled with quantitative PCR (RT-qPCR) (**Figure 5d**). For this analysis, we chose 5 of the top 10 downregulated transcripts (highlighted in **Figure 5c**). In this approach, total RNA, without rRNA depletion, was independently purified from that used in the RNA-seq analysis. For reverse transcription, random priming was used to avoid potential bias from changes in poly-adenylation. First, we confirmed the degree of NOCT over-expression and observed a 2.9-fold increase, consistent with the 3.1-fold increase measured by RNA-seq. Next, we measured the change in expression of the mRNAs encoding *DLGAP1, RORB, TMEFF2, PTPRZ1*, and *FRRS1L* mRNAs. DLGAP1 (Disks Large-Associated Protein 1) is a neuron-specific component of the post-synaptic scaffold which are linked to glutamate receptors (33). RORB (Retinoic acid receptor-related Orphan Receptor β) is a member of the nuclear hormone receptor family of DNA binding transcription factors which regulates brain development and bone formation (34,35). TMEFF2 (Trans-Membrane protein with EGF like and two Follistatin like domains 2) is a transmembrane protein linked to multiple cancers (36). PTPRZ1 (Protein Tyrosine Phosphatase Receptor type Z1) is a brain specific protein that regulates neuronal development and cell migration (37). FRRS1L (Ferric Chelate Reductase 1 Like) is part of the neuronally expressed AMPA receptor, mutations of which are linked to infantile epileptic encephalopathy, and deletions cause intellectual disability (38,39). For each transcript, the fold-change measurements made by RT-qPCR were concordant to those measured by RNA-seq (**Figure 5d**). We also measured the levels of three mRNAs that were not affected by NOCT over-expression in the RNA-seq dataset; the RT-qPCR analysis confirmed that beta 2-microglobulin (B2M), beta-actin (ACTB), and TATA binding protein (TBP) mRNAs are unaffected by NOCT over-expression. Thus, two independent approaches corroborate that NOCT significantly changes expression of specific genes in a reproducible manner. Taken together, our data indicate that cytoplasmic NOCT selectively regulates gene expression.

To assess the potential effect of NOCT on biological functions and processes, we performed information-theoretic pathway analysis using iPAGE (40). Gene ontology terms that are either significantly over- or under-represented in the set of genes affected by NOCT Δ(2-15)-3F in our RNA-seq dataset are plotted in **Figure 6**. We focus on GO terms that are enriched in the downregulated genes because these are the most likely to be repressed by NOCT. This category includes significantly enriched terms such as homophilic cell adhesion via plasma membrane adhesion molecules, transmembrane receptor tyrosine kinase activity, and negative regulation of osteoblast differentiation. The latter category is intriguing because NOCT was reported to inhibit osteoblastogenesis and negatively regulate bone density in mice (9,10,41). **Supporting Information Figure S2** provides gene level data for these over-represented categories. We also observed enrichment of multiple terms associated with neuronal functions including ionotropic glutamate receptor activity, neurotransmitter secretion, and voltage-gated potassium channel activity. The latter group includes transcripts from the genes *KCNQ5, KCNA4, KCNV1, KCND3, KCNT2*, and *KCNQ2*, which are voltage-gated potassium channels that are expressed in the brain and are linked to neurological disorders including intellectual disability, microcephaly, seizures, movement disorders, and spinocerebellar ataxia (42,43). These results provide new evidence that NOCT is capable of downregulating neuronal transcripts and suggests a new biological role for cytoplasmic NOCT. Given the range of tissue types which show native NOCT expression, it is likely that NOCT orchestrates different processes in different cell types.

**Figure 6.**
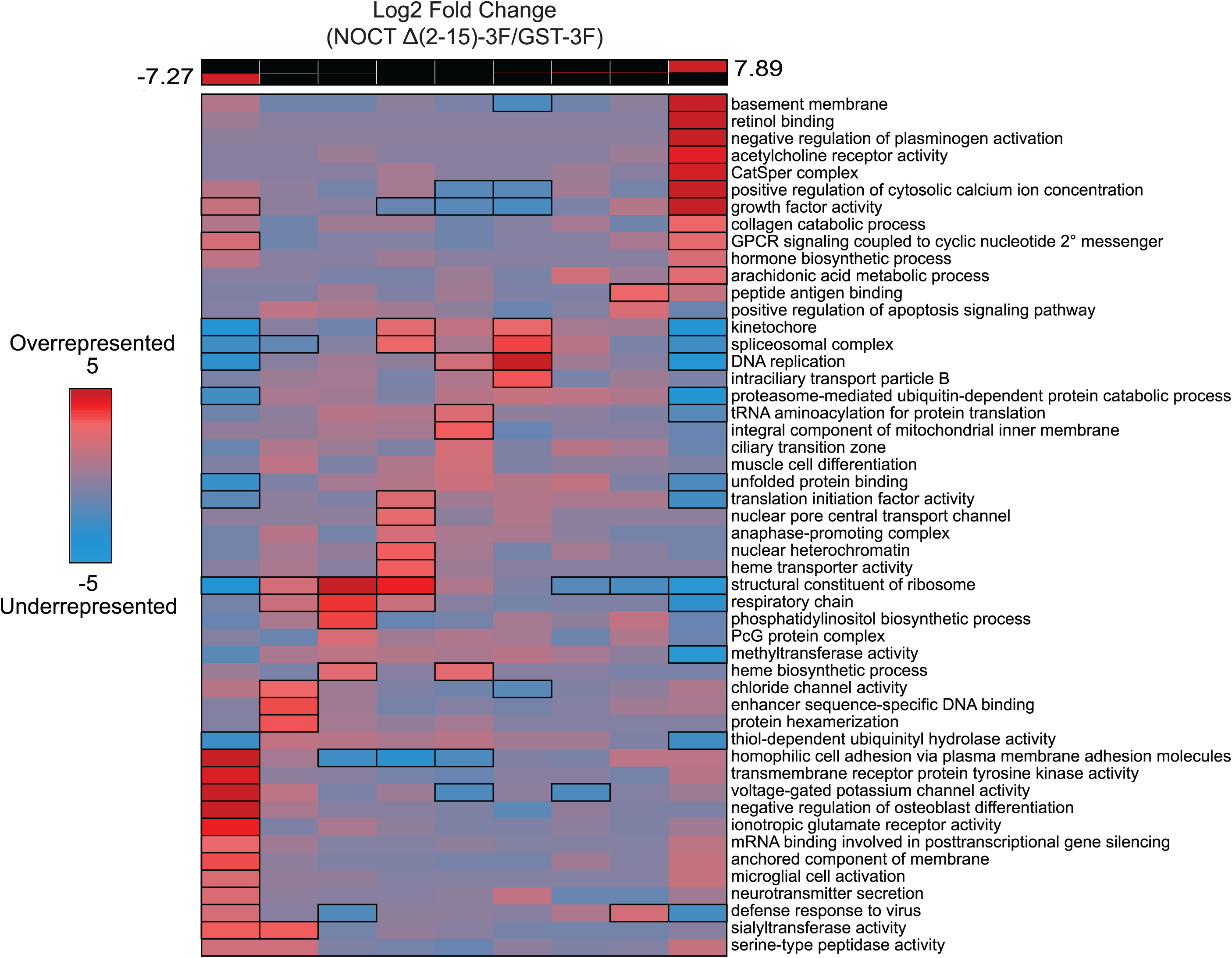
Pathway enrichment analysis of gene expression changes in response to NOCT Δ(2-15)-3F. RNA-seq data was analyzed using iPAGE to identify pathways with significant mutual information using gene specific log2 fold changes estimated by DESeq2 without shrinkage. The upper panel shows the discretized bins of the gene-specific metric as ranges of expression changes within each bin in the context of the overall range (red boxes in black bins). The leftmost bins contain genes with the lowest log2 fold change metrics, with the rightmost bins containing the highest. The colors of tiles correspond to the over-(red) or under-represented (blue) gene ontology (GO) terms, with color intensity quantified as log10 of the GO term enrichment p-values. Therefore, more saturated red or blue colors indicate higher significance. Bold borders of tiles indicate significant enrichments within an expression bin (p < 0.05 after Bonferroni correction across the row); note that, however, all displayed GO terms have significant mutual information given the overall gene expression change profile (as assessed by the default series of tests used by iPAGE). The RNA-seq data and statistics used in this analysis are described in Supporting Information Table S1.

### NOCT affects cellular levels of nicotinamide adenine dinucleotide cofactors

A recent study reported that NOCT can dephosphorylate the metabolites NADP^+^ and NADPH in biochemical assays (22). We therefore sought to analyze the effect of differentially localized NOCT on these metabolites in live cells. We predicted that NOCT phosphatase activity may decrease levels of NADP(H) and increase the levels of cellular NAD(H). In order to assess NOCT-mediated changes in metabolite levels, we first created HEK293 derived cell lines with stably integrated, doxycycline (Dox)-inducible NOCT, including mitochondrial localized NOCT (1-431)-3F or cytoplasmic NOCT Δ(2-15)-3F. Three independent clonal cell lines were created for each construct, and matched wild type cell lines were used as negative controls. Treatment with Dox for 72 hours led to induction of NOCT Δ(2-15)-3F protein (**Figure 7a**) with the anticipated ∼58 kDa apparent molecular weight. Likewise, Dox-induced NOCT (1-431)-3F protein was specifically expressed and processed to the ∼44 kDa apparent molecular weight (**Figure 7b**). We also measured the level of induction of each NOCT construct by RT-qPCR and observed 14-fold and 4.2-fold induction of NOCT Δ(2-15)-3F and NOCT (1-431)-3F, respectively (**Figure 7c**). Next, we measured the effect of induced NOCT on relative levels of NAD^+^, NADH, NADP^+^, and NADPH levels in each cell line using luminescence-coupled assays. To assess reproducibility, we performed three independent experiments, each of which included 3 biological replicates for each test condition. The resulting data were analyzed by calculating the fold change of each metabolite in each cell line between the Dox-induced state relative to the non-induced, vehicle-treated state. We applied a Bayesian statistical analysis of the resulting data to assess reproducibility and significance. Mean log2 fold change values are plotted in **Figure 7d**, along with 95% credible intervals and significance calling.

**Figure 7.**
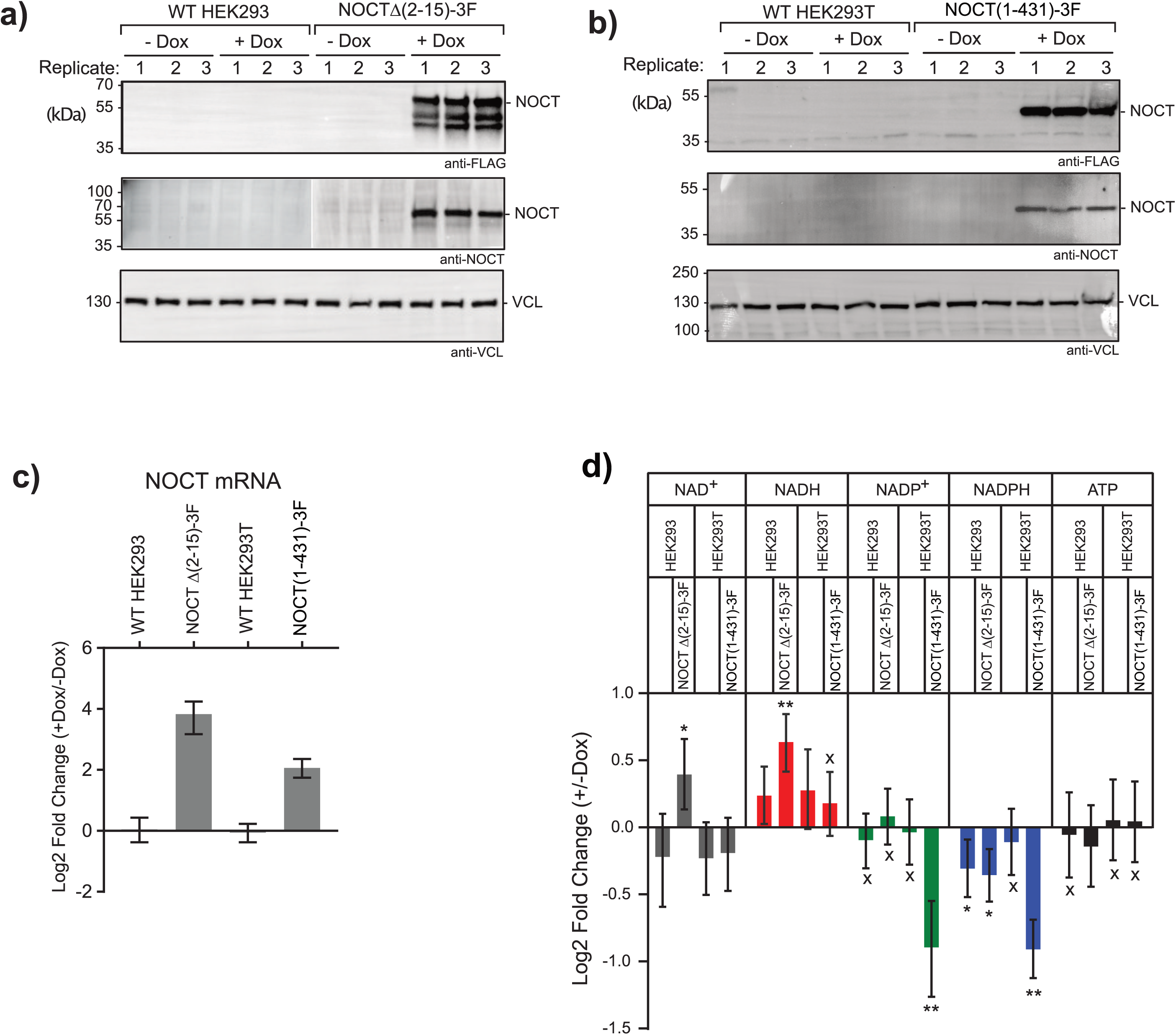
Effect of NOCT on cellular levels of nicotinamide adenine dinucleotide cofactors and ATP. **a)** Dox-inducible expression of NOCT Δ(2-15)-3F was confirmed in three separate pools of stably transduced HEK293 cells by performing anti-FLAG and guinea pig anti-NOCT western blotting. NOCT Δ(2-15)-3F protein migrated with an apparent molecular weight of 58 kDa, consistent with unprocessed NOCT. Wild type (WT) HEK293 cells served as negative controls. Cells were treated with 1 µg/mL Dox (+ Dox) for 72 hours or the DMSO vehicle (-Dox). Equal mass of each cell lysate was loaded into each well and vinculin (VCL) was detected by western blot to assess equivalent loading of lanes. **b)** As in a), but using HEK293T cells with Dox inducible NOCT (1-431)-3F that was introduced by the Flp-In T-REx system. NOCT (1-431)-3F protein migrated with a molecular weight of 44 kDa, consistent mitochondrial protease processing. Wild type HEK293T cells served as negative controls. **c)** Over-expression of the stably expressed Dox-inducible NOCT constructs in the cell lines used in a) and b) were quantitated at the mRNA level using RT-qPCR. Each bar in the graph represents n=9 from three replicate experiments with three biological replicates each. Log2 fold change values were calculated for NOCT mRNA in the presence of Dox relative to without Dox. Mean log2 fold change values are plotted along with 95% credible intervals. Data and statistics are reported in Supporting Information Table S2. **d)** NOCT Δ(2-15)-3F significantly increased levels of NAD+ and NADH, while NOCT (1-431)-3F significantly decreased levels of NADP+ and NADPH. The Dox-inducible and control cell lines were treated with either DMSO or 1 µg/mL Dox for 72 hours. The levels of NAD+, NADH, NADP+, NADPH, and ATP were then measured in equal numbers of cells for each condition. Mean log2 fold change in each metabolite was calculated in the Dox-induced condition relative to the uninduced condition. Mean log2 fold change values are plotted for each measurement, along with 95% credible intervals. Measurements were derived from three replicate experiments, each of which contained three biological replicates. For significance calling, a ‘*’ denotes a posterior probability >0.95 that the difference relative to the negative control is in the indicated direction. The ‘**’ indicates a posterior probability of >0.95 that the indicated difference is at least 1.3-fold. An ‘x’ marks a posterior probability >0.95 that the indicated difference is no more than 1.3-fold in either direction. Data and statistical values are reported in Supporting Information Table S2.

The resulting data demonstrate that NOCT Δ(2-15)-3F caused a significant increase in the levels of NAD^+^ and NADH. These results suggest that cytoplasmic NOCT can dephosphorylate NADP(H) to increase the level of NAD^+^/NADH. However, NOCT Δ(2-15)-3F did not significantly reduce the pool of either NADP^+^ or NADPH relative to control cells. Induced NOCT (1-431)-3F significantly reduced the levels of NADP^+^ and NADPH, indicating that when localized to the mitochondria, NOCT can dephosphorylate NADP^+^ and NADPH. The pool of NAD^+^ and NADH was not detectably changed by mitochondrial NOCT. Together, these results provide evidence that NOCT can alter nicotinamide adenine dinucleotide levels in a manner dependent on its intracellular localization. We also investigated the potential impact of NOCT on overall ATP levels and observed that neither cytoplasmic NOCT Δ(2-15)-3F nor mitochondrial NOCT (1-431)-3F affected ATP levels. Therefore, NOCT over-expression does not appear to disrupt cellular bioenergetics, nucleotide biogenesis, or cell viability.

## Discussion

The results of this research provide important new insights into the molecular functions, expression pattern, and intracellular localization of human NOCT and its impact on metabolism of mRNAs and nicotinamide adenine dinucleotide cofactors. We found that NOCT protein is expressed and differentially processed in a cell-type and tissue-specific manner. The processing is consistent with mitochondrial targeting of NOCT. Indeed, we showed that NOCT can be localized into mitochondria, dependent on an N-terminal mitochondrial targeting sequence. In corroboration, two new studies also detected over-expressed NOCT in mitochondria (22,23).

Human *NOCT* mRNA is broadly expressed but is most abundant in tissues such as adipose, breast, lung, liver, skeletal muscle, and kidney (**Supplementary Figure 1**). We have additionally observed that NOCT protein is abundant in liver, kidney, skeletal muscle, and brain. Interestingly, kidney and liver exclusively express the unprocessed ∼55 kDa form of NOCT (which is presumably retained in the cytoplasm), whereas skeletal muscle expressed only the ∼40 kDa form and brain produced both forms (**Figure 4**). These observations indicate that NOCT processing and localization are regulated. The mechanism of such regulation remains unknown and should be the focus of future studies. While several other examples of differential cytoplasmic-mitochondrial localization have been reported, those cases involve utilization of distinct translation initiation sites to produce protein isoforms that either do or do not possess an MTS (44-46). Our data indicate that NOCT translation initiates at M1 and thus differential localization of NOCT is controlled through regulation of MTS-mediated import and processing. In one reported case, both types of mechanisms appear to operate: differential usage of translation initiation sites can produce the CNP2 protein isoform that has an extended N-terminal containing an MTS, and subsequent phosphorylation of the CNP2 MTS prevents its mitochondrial localization (46). While phosphorylation sites have been mapped in NOCT, they do not occur within the MTS, though it is noteworthy that the NOCT MTS has two serine residues (S4 and S10). Future research will be necessary to determine the precise mechanism that controls NOCT processing and localization in different tissues.

Differential localization of NOCT to the cytoplasm or mitochondria has important implications for the repertoire of substrates it may act on. Cytoplasmic NOCT would target nuclear-encoded mRNAs, which is consistent with the changes in mRNA abundance observed when NOCT expression is perturbed (15-17). Our results provide evidence that cytoplasmic NOCT specifically regulates a network of hundreds of mRNAs in HEK293 cells. It is noteworthy that the vast majority of mitochondrial proteins are nuclear-encoded, and overall in our RNA-seq analysis, we saw little or no change in their expression levels (**Supporting Information Figure S3** and **Table S1**). In contrast, localization of NOCT to the mitochondria would restrict the pool of potential targets to 13 mRNAs, 2 rRNAs and 22 tRNAs (47). Mitochondrial transcripts encode a subset of the subunits of the electron transport chain, and their regulation should affect metabolism and cellular energy states (48). While the impact of mitochondrial NOCT on mitochondrial RNAs remains to be fully explored, we did not detect altered cellular bioenergetics as assessed by measuring ATP levels, when either mitochondrial or cytoplasmic NOCT was over-expressed, indicating that mitochondrial gene expression was not altered significantly. Moreover, a recent report analyzed the effect of knockout of NOCT in mouse brown adipose, and observed little to no effect on mitochondrial gene expression, though a subset of mitochondrial encoded mRNAs showed a modest decrease in expression during cold adaptation (18).

To gain insight into the genes and biological functions that NOCT can control, we analyzed the effects of cytoplasmic NOCT on expression of nuclear-encoded mRNA transcripts in HEK293 cells. HEK293 cells were advantageous in that they express a low level of endogenous NOCT, and thus our over-expression approach mimics the induction of NOCT in response to circadian inputs, mitogens, lipids, or other stimuli (4,8,11,49,50). Some 490 transcripts were significantly affected, and expression of 235 of those transcripts decreased significantly, consistent with NOCT-mediated repression. Multiple gene ontology terms were enriched in the downregulated genes, notably including genes involved in cell adhesion, negative regulation of osteoblast differentiation, receptor tyrosine kinase activity, and neurological functions (voltage-gated potassium channel activity, ionotropic glutamate receptor activity, microglial cell activation, neurotransmitter secretion) (**Figure 8a**). Relevant to this trend, endogenous NOCT is expressed in both human and mouse brain tissues (**Figure 4**). It is noteworthy that previous studies examined NOCT-mediated changes in gene expression in mouse adipose and liver tissue, but not neuronal cell types. While the human HEK293 cells used in our analysis were originally derived from embryonic kidney, they exhibit kidney, neuronal, and adrenal phenotypes, which likely facilitated our detection of affected neuronal genes (31,32). Based on our observations, the potential regulatory roles for NOCT in neurological functions should be further investigated using tools such as brain-specific knockout models and examination of neurological functions and phenotypes.

**Figure 8.**
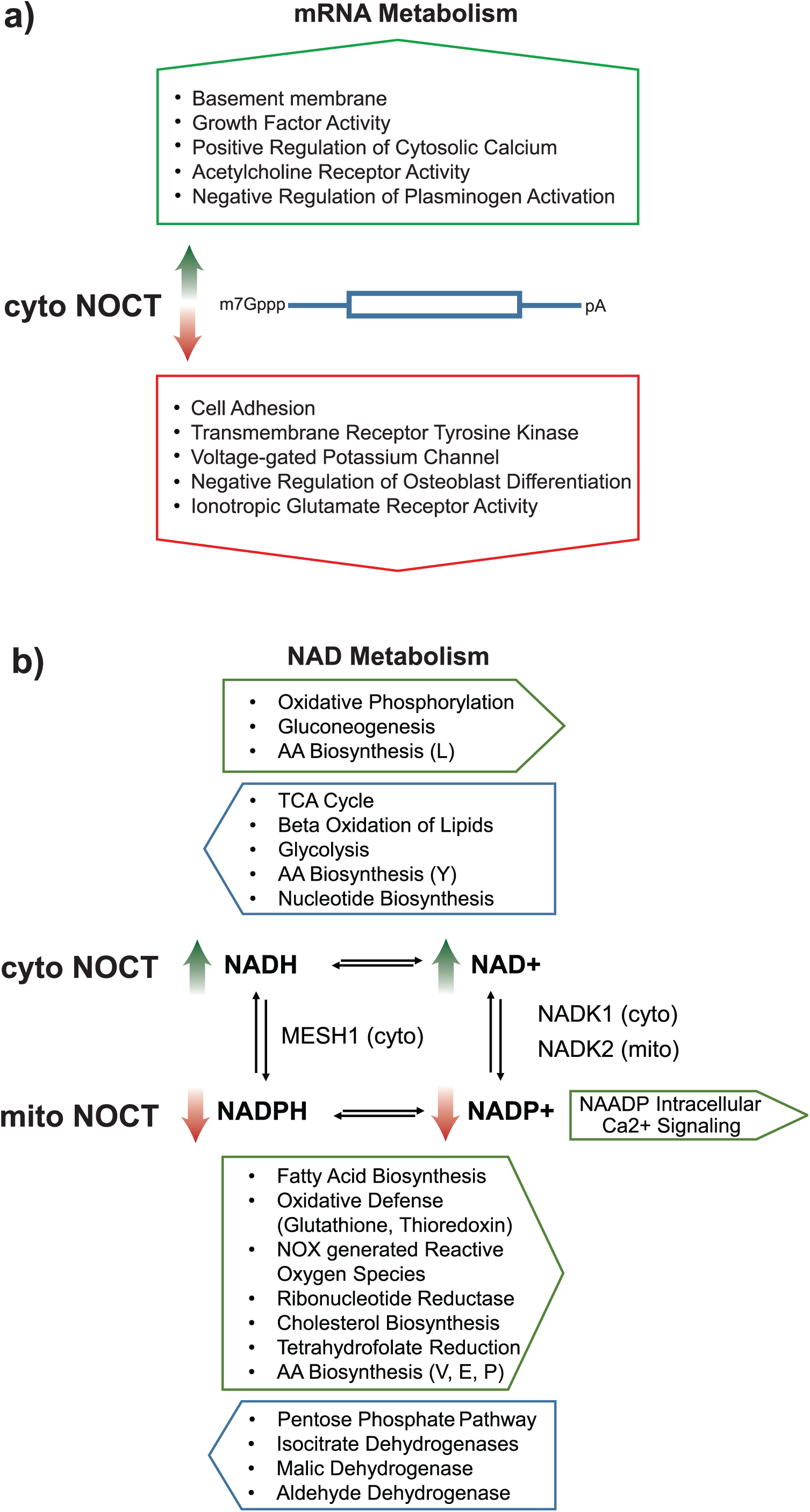
Summary of NOCT-mediated effects on mRNA and NAD metabolism. **a)** Cytoplasmic, unprocessed NOCT changed the expression of mRNAs from specific genes, including those enriched with the indicated gene ontologies. The green arrow indicates genes whose levels were increased by cytoplasmic NOCT, whereas the red arrow indicates genes whose levels were decreased. **b)** Diagram of NAD cofactor metabolism and the effect of over-expressed forms of NOCT. NAD cofactors are metabolized by a variety of anabolic and catabolic pathways, indicated above and below, which interconvert NAD cofactors between redox states. NADP+ can also be converted to the signaling molecule NAADP. In addition, NAD cofactors are interconverted between phosphorylation states, catalyzed by at least two kinases (NADK1 and NADK2) and phosphatases (MESH1 and NOCT). These kinases exhibit specific subcellular localization between cytoplasm (cyto) and mitochondria (mito), as indicated. Green arrows indicate the observed effect of cytoplasmic NOCT on increased NADH and NAD+ levels. Red arrows indicate the observed effect of mitochondrial NOCT on decreased levels of NADPH and NADP+.

The relationship between the effects of NOCT on transcript levels to the reported lipid metabolism phenotypes remains obscure. Knockout of NOCT in mice resulted in resistance to high fat diet induced obesity, coupled with reduced uptake, transport, and storage of fat. Further, knockout mice had reduced adipose tissue (4,8,9). We did not observe significant changes in lipid metabolism pathways in our dataset.

NOCT was also reported to negatively regulate osteoblastogenesis and knockout of NOCT resulted in an increase in bone density (10,41,51). Interestingly, we identified RORB mRNA as a NOCT-repressed target. RORB is a regulator of osteoblastogenesis and affects bone density and resorption during aging (35). This information suggests that RORB could be an important target of NOCT in bone development and maintenance.

Collectively, our RNA-seq data provide further evidence, coupled with previous studies, demonstrating a role of NOCT in mRNA metabolism (1,15-17). The observation that NOCT reduces the abundance of hundreds of mRNAs when over-expressed is consistent with its proposed function as an exoribonuclease. Though pure NOCT protein does not exhibit deadenylase activity in biochemical assays, in vivo it can negatively regulate mRNA levels (1,12). Further support for the proposed exoribonuclease activity comes from tethered function assays, wherein NOCT reduced RNA level when directed to a reporter mRNA. This activity was diminished by mutations of certain conserved residues in the active site of NOCT. Moreover, incorporation of a stable triple helical RNA structure into the 3′ end of the reporter mRNA abrogated NOCT-mediated repression (1). Additional support for repression of mRNAs by NOCT comes from analysis of the effects of deletion or RNAi-mediated depletion of NOCT, which caused upregulation of a subset of mRNAs in several model systems, including mouse liver cells and cultured mouse adipocytes (15-17). The fact that NOCT affects a subset of transcripts indicates that a mechanism conferring transcript specificity exists but remains unknown. Other RNA-degrading enzymes can be directed to their target RNAs through sequence specific recognition of cis-acting RNA elements, often conferred through partnership with RNA-binding proteins (7). We were unable to identify enrichment of cis-regulatory elements in the NOCT-regulated transcripts in our dataset. Future studies are necessary to determine how NOCT regulates these RNAs and to identify the protein partners that may regulate its putative exoribonuclease activity and substrate specificity.

A recent study reported that purified NOCT can dephosphorylate the cofactors NADP^+^ and NADPH in vitro; therefore, we explored the potential impact of NOCT on cellular pools of nicotinamide adenine dinucleotide cofactors (22). We observed that over-expression of cytoplasmic NOCT increased the cellular pool of NADH and NAD^+^ levels whereas mitochondrial NOCT decreased the level of NADP^+^ and NADPH (**Figure 8b**). Overall, these observations are consistent with the reported NADP(H) phosphatase activity of NOCT. The implications are intriguing but complex. NAD cofactors exhibit high flux between redox states through crucial roles in many pathways, including both biosynthetic and catabolic pathways, cellular bioenergetics, protection from and generation of reactive oxygen species, and signal transduction (summarized in **Figure 8b**) (52-54). NAD cofactors also interchange between cytoplasmic and mitochondrial compartments, either indirectly through shuttle pathways or — as indicated by recent evidence — directly (53,55-58). Most relevant to NOCT, NAD is also interconverted between phosphorylation states, mediated by specific cytoplasmic and mitochondrial NAD kinases, NADK1 and NADK2, respectively (**Figure 8b**) (59,60). Additionally, another cytoplasmic NADPH phosphatase, MESH1/HDDC3, has been described (61). Interestingly, the expression pattern of MESH1 across human tissues is broader and higher level than NOCT (**Supporting Information Figure S1**). It is noteworthy that we explored the possibility that NOCT might alter expression of these key NAD interconverting enzymes; while NADK1, NADK2, and MESH1 are well-expressed in our RNA-seq data, none were significantly affected by over-expressed cytoplasmic NOCT (**Supporting Information Table S1**). Our data, coupled with that of Estrella et al., indicate that the NADP(H) phosphatase activity of NOCT contributes to NAD cofactor interconversions. The differential localization of NOCT to cytoplasm or mitochondria suggests that it may alter NAD-linked metabolism in a cell type and tissue type specific manner.

A broad implication of the NADP(H) phosphatase activity of NOCT is that it could alter cellular metabolism. Dephosphorylation of NADP(H) would increase cellular NAD(H) pools, and generally promote catabolic pathways including glycolysis, beta-oxidation of fatty acids, and oxidative phosphorylation. Such changes would be anticipated to promote ATP production, though we did not detect such an effect. It is also worth considering how NOCT NADP(H) phosphatase activity may relate to the reported lipid metabolism and storage defects observed in the knockout mice. Loss of NOCT activity should increase the pool of NADP(H), and thereby promote biosynthetic pathways including synthesis of lipids. This prediction is at odds with the observed loss of adipose tissue and defects in lipid update, transport, and storage observed in the NOCT knockout mice (4,8). Clearly more research is necessary to determine whether the NADP(H) phosphatase activity of NOCT is relevant to the reported resistance to fat-induced obesity phenotype, or whether the exoribonuclease function of NOCT is more important for this phenotype.

The emerging picture indicates that NOCT may be a dual function enzyme that controls RNA and NADP(H) metabolism. Thus, it joins the growing ranks of “moonlighting” metabolic enzymes that fit the REM (RNA-Enzyme-Metabolite) hypothesis (62). The classic REM example is aconitase, an iron-sulfur cluster, citric acid cycle enzyme that also binds and regulates transcripts involved in iron homeostasis (63-66). Recent global surveys have identified more examples of moonlighting RBPs, suggesting that many of these dual function enzymes provide a level of translational control in response to metabolic states (62). Interestingly, six NAD cofactor utilizing enzymes have been found to also bind to mRNAs. One example is Glyceraldehyde-3-phosphate dehydrogenase (GAPDH), which binds to AU-rich RNA elements of specific mRNA transcripts within the same binding site that binds NAD^+^ (67-69). Future work on NOCT will be crucial for determining the contributions of the NOCT NADP(H) phosphatase and mRNA repression activities and differential localization to the control of gene expression and metabolism.

## Experimental Procedures

### Plasmids

The pFC3F plasmid backbone was generated from the pF5A vector (Promega) to generate C-terminal 3x FLAG tag fusions. Human NOCT (1-431) or *Schistosoma japonicum* GST coding sequences were inserted into pFC3F using the Flexi Cloning System (Promega). To generate NOCT Δ(2-15)-3F, residues 2-15 were removed using inverse PCR. NOCT N-terminal point mutants were generated from pCR4 TOPO NOCT (Open Biosystems) and Quikchange site directed mutagenesis PCR (Agilent) to generate NOCT(1-431) M1X, NOCT (1-431) M67G, and NOCT (1-431) R15M. NOCT Δ(2-15) and NOCT Δ(2-15) M67G were generated from pCR4 vectors using inverse PCR. All NOCT inserts generated in pCR4 TOPO were then transferred to pF5A using the Flexi Cloning System (Promega). To generate NOCT (1-86) EGFP, the EGFP ORF was PCR amplified from the pEGFPN1 vector (Clontech) with appended 5′ AfeI and 3′ EcoRI restriction sites. pF5A NOCT(1-431) was digested with AfeI and EcoRI to remove NOCT (87-431), which was replaced by the EGFP ORF using T4 DNA ligase (NEB). We note that the initiator methionine codon of the EGFP ORF was not included in this construct. NOCT Δ(2-15)-3F pCW57.1 was generated using the pCW57.1 (Addgene, #41393) plasmid backbone. NOCT Δ(2-15)-3F was PCR amplified from NOCT Δ(2-15)-3F pFC3F and inserted into pCW57.1 using Gibson Assembly using the NEBuilder HiFi Assembly Master Mix (NEB, E5520). NOCT (1-431)-3F pcDNA5/FRT/TO was generated through PCR amplification of NOCT (1-431) cDNA (Source Biosciences I.M.A.G.E. clone 8322546) with appended 5′ BamHI and 3′ NotI restriction sites. The NOCT PCR insert and pcDNA5/FRT/TO were digested with BamHI and NotI restriction enzymes (NEB) before ligation with T4 DNA ligase (NEB). For all constructs, positive clones were confirmed by DNA sequencing.

### Generation of α-NOCT antibodies

The rabbit α-NOCT polyclonal antibody was produced in rabbits inoculated with MBP-NOCT (1-431) (Thermo Fisher custom antibody services). This MBP-NOCT antigen (1-431) was recombinantly expressed in BL21 DE3 cells and purified over amylose resin. Crude serum was tested for reactivity against HEK293 cell extracts overexpressing HaloTag-NOCT (1-431). NOCT antibodies were then purified from the verified serum by ammonium sulfate fractionation and then antigen affinity purification using recombinantly expressed HaloTag-NOCT (1-431) immobilized on HaloLink resin (Promega). The serum was purified over the immobilized NOCT column before elution with 5 M NaI and 1 mM ammonium thiocyanate. Antibody was subsequently dialyzed into 1x PBS (137 mM NaCl, 27 mM KCl, 10 mM Na_2_HPO_4_, and 18 mM KH_2_PO_4_) for 1 hour at 4 °C. Dialysis was repeated two additional times with the final dialysis step performed overnight. Insoluble material was removed by centrifugation at 18,000 x g and the antibody was concentrated to 1 mg/mL using a Centricon spin concentrator (Millipore Sigma). Purified polyclonal antibody was verified for reactivity against recombinant purified NOCT (64-431) (purified as described in Abshire et al, 2018), as shown in **Figure 1** (1).

The guinea pig α-NOCT polyclonal antibody, used in **Figures 2 and 7**, was produced in guinea pigs inoculated with Strep(II)-Sumo-NOCT (64-431) (Cocalico custom antibody services). Briefly, Strep(II)-Sumo-NOCT (64-431) was recombinantly expressed in BL21 DE3 Rosetta2 STAR cells and purified over Strep-Tactin Superflow Plus resin (Qiagen). Crude serum was verified for reactivity against recombinant NOCT (64-431) and HEK293 cell extracts from HEK293 cells stably overexpressing NOCT (1-431)-3F. Antibody from verified serum was purified using Protein A agarose (Pierce) and the antibody fraction was eluted using 0.1 M glycine and 20% glycerol. Eluates were neutralized with 0.15 M Tris, pH 8 and the α-NOCT antibody was antigen affinity purified with NOCT (64-431) immobilized on Immobilon-P PVDF membrane (Millipore). Guinea pig α-NOCT was eluted from the membrane with 0.1 M glycine and 20% glycerol. Eluates were neutralized with 0.15 M Tris, pH 8 before the antibody was concentrated to ∼0.1 mg/mL using a Centricon spin concentrator. Purified polyclonal antibody was verified for reactivity against recombinant purified NOCT (64-431) and lysates from HEK293 cells stably overexpressing NOCT (1-431)-3F.

### Analysis of differential NOCT processing by western blotting

HEK293 cells were cultured in DMEM (Fisher), 1% Pen/Strep/Glutamine (Fisher) and 10% FBS (Thermo) at 37 °C and 5% CO_2_. Cells were transfected with NOCT N-terminal mutant constructs using Fugene HD (Promega) in a ratio of 3 µL transfection reagent per 1 µg DNA. HCT116 cells were cultured as described for HEK293 cells but in McCoy’s 5A medium (Fisher), 1% Pen/Strep/Glutamine and 10% FBS. Cells were transfected using Fugene HD (Promega) in a ratio of 4 µL transfection reagent per 1 µg DNA. Both HEK293 and HCT116 cells were harvested and resuspended in modified radio-immunoprecipitation assay buffer (mRIPA buffer, 10 mM Tris (pH 8.0), 140 mM NaCl, 1 mM Diaminoethane-tetraacetic acid (EDTA), 0.5 mM Triethylene glycol diamine tetraacetic acid (EGTA), 1% (v/v) Triton X-100, 0.1% (w/v) sodium deoxycholate, and 0.1% (w/v) sodium dodecyl sulfate) for lysis containing complete protease inhibitor cocktail (Roche). Lysates were homogenized using a handheld cell disruptor and sample concentrations were determined using a DC Lowry protein assay (BioRad). For cell lysates, equal masses of protein samples (20 µg) were resolved using SDS PAGE and then transferred to Immobilon-P PVDF membrane (Millipore). Recombinant NOCT (64-431) (6 ng) was used as a positive control for NOCT and was expressed and purified as previously described (1). Blots were probed using the following primary antibodies: α-NOCT (derived from rabbit serum, detecting NOCT MTS mutants expressed from pF5A), α-EGFP (detecting EGFP and NOCT (1-86)-EGFP, Clontech clone JL-8), and α-FLAG (detecting C-terminal 3x FLAG-tagged constructs, Sigma). Blots were probed using HRP-conjugated Secondary Antibody (Sigma) and Immobilon Western HRP Chemiluminescent substrate (Millipore) before imaging on a Chemidoc Touch (BioRad). As loading controls, either Vinculin (VCL) or GAPDH were detected on the same blots using α-Vinculin antibody (Thermo Fisher) or α-GAPDH antibody (Ambion).

### Analysis of Human and Mouse Tissue blots

Pre-transferred blots of tissue extract panels from human and mouse were purchased from Novus Biologicals (Insta-blot NBP2-31378 and NBP2-20111). According to the manufacturer, each blot contains 50 µg of total protein from the indicated tissue type. Tissues were from donors with no known disease and proteins were extracted using lysis buffer (10 mM Tris, pH 8.0, 130 mM NaCl, 1% Triton X-100, 10 mM NaF, 10 mM NaPi, 10 mM NaPPi) with protease inhibitor cocktail. Blots were hydrated and blocked before probing with the α-NOCT antibody derived from rabbit serum. Blots were probed using HRP conjugated goat α-rabbit IgG secondary antibody (Sigma) and Pierce ECL Western blotting substrate before imaging on a Chemidoc Touch (BioRad).

### Generation of Cell lines

All parental cell lines were purchased from the American Type Culture Collection, unless otherwise noted. To produce stably transfected HEK293 cell lines (ATCC), cells were transfected with a 1:10 molar ratio of pEF6 HisA (Invitrogen) to either GST-3F pFC3F or NOCT Δ(2-15)-3F pFC3F using Fugene HD. 5 µg/mL Blasticidin S (Thermo Fisher) was used to select Blasticidin resistant populations that were then subcloned using the dilution method. Blasticidin resistant clones expressing 3x FLAG-tagged fusion proteins for each construct were identified by western blotting using α-FLAG. Briefly, cells were harvested in 1x PBS, pelleted, and frozen. Prior to use, cell pellets were resuspended in mRIPA supplemented with protease inhibitor (Millipore Sigma) for lysis. Lysates were homogenized using a handheld cell disruptor and sample concentrations were determined using the BioRad DC Lowry assay. Each sample (10 μg total protein) was resolved on 4-20% TGX gradient gels (BioRad) and then transferred onto Immobilon-P PVDF membrane. Blots were probed using α-FLAG (Sigma) before detection using HRP-conjugated Secondary Antibody (Sigma) and ECL Western Blotting substrate (Pierce) before imaging on a Chemidoc Touch (BioRad).

To generate HEK293 cells expressing NOCT Δ(2-15)-3F under the control of the tet operator, lentivirus was produced in HEK293FT cells transfected with NOCT Δ(2-15)-3F pCW57.1 and the pRSV-Rev (Addgene #12253), pMD2.G (Addgene #12259), and pMDLg/pRRE (Addgene #12251) packaging vectors. HEK293 cells were transduced with 400 μL of viral supernatant and 6 μg/mL polybrene in 1 mL final culturing volume per well in a 6-well plate. The population of treated HEK293 cells were selected for cells with genomic integration with 1 μg/mL puromycin. A population of puromycin resistant cells were treated with 1 µg/mL Doxycycline (Sigma) for 48 hours. To verify expression of NOCT Δ(2-15)-3F the transduced and selected populations of HEK293 cells were harvested, pelleted, and lysed as described above. Expression of NOCT Δ(2-15)-3F was verified by western blotting with α-FLAG (Sigma).

To produce tet-inducible, Flp-In NOCT (1-431)-3F HEK293T cell lines, parental Flp-In HEK293T cells (Invitrogen) were transfected with NOCT (1-431)-3F pcDNA5/FRT/TO and pOG44 using Lipofectamine 3000 (Invitrogen) according to the manufacturer’s instructions. Cells with genomic integration were selected for using 100 μg/ml Hygromycin (Gibco) and 10 μg/ml Blasticidin (Gibco). Single colonies were generated from the antibiotic resistant population and verified using α-FLAG Western blotting.

### Immunofluorescence

Human 143B osteosarcoma (HOS) cells were cultured for use in immunocytochemistry experiments in DMEM supplemented with 10% (v/v) FBS, 2 mM GlutaMax (Gibco) and 1× Penicillin/Streptomycin at 37°C under 5% CO_2_. For each construct, 2 μg of plasmid was transfected into HOS cells at 40% confluency using Lipofectamine 3000 transfection reagent (Thermo Fisher) according to manufacturer’s instructions. After 24 hours, cells were split onto microscope slides, and at 48 hours post-transfection, cells were incubated for 35 minutes with 200 nM MitoTracker Red CMXRos (ThermoFisher) at 37°C. Cells were washed 1x with pre-warmed DMEM, and once with D-PBS before fixing in PBS + 4% formaldehyde for 15 minutes at room temperature. Cell permeabilization was performed by incubating cells in PBS + 5% FBS (PBSS) supplemented with 1% Triton X-100 for 5 minutes at RT. Cells were washed twice with 1x PBSS (without Triton X-100) and blocked with PBSS for 1 hour. To visualize 3x FLAG-tagged proteins, cells were incubated with 1:200 mouse anti-FLAG M2 antibody (Sigma-Aldrich, F3165) for 2 hours at RT, followed by triplicate 5 minute washes with PBSS. Cells were then incubated with 1:1000 Goat anti-mouse Alexa Fluor Plus 488 (ThermoFisher, A32723) for 1 hour at RT before slides were washed 3x with PBSS for 5 minutes each. DAPI staining (300 nM, Sigma-Aldrich) was performed during the first of the final washes. Immunofluorescence images were captured using a Zeiss LSM800 confocal microscope and Fiji software was used to produce the merged images.

### Subcellular fractionation of cells

For subcellular fractionation of NOCT (1-431)-3F HEK293T cells, mitochondrial isolation was performed as previously described, with the omission of the sucrose gradient step (70). Mitochondrial pellets were gently resuspended in MSH buffer (210 mM mannitol, 175 mM sucrose, 20 mM DTT, 0.2 mM PMSF, and 1x protease inhibitor cocktail (Roche)). The protein concentration of the total cellular, cytoplasmic, and mitochondrial fractions was measured using the DC Lowry protein assay (BioRad) and split into 3 equal aliquots. Mitochondrial fractions were subjected to no treatment, treatment with Proteinase K (20 μg/4 mg mitochondrial protein, Thermo Fisher), or treatment with Proteinase K + 1% Triton X-100 to expose intramitochondrial proteins to protease activity. Equal volumes of treated mitochondrial fractions, and cytoplasmic and nuclear fractions with protein concentrations equal to the untreated mitochondrial fraction were resolved via SDS PAGE followed by Western blotting as described above. Immunoblotting was performed using the following antibodies: Goat anti-DDDDK (FLAG), (1:5000 Abcam, ab1257), Guinea pig anti-NOCT (described above), Rabbit anti-uL4m antibody (1:2000, Sigma, HPA051261), Rabbit anti-uS15m antibody (1:2000 ProteinTech Group, 17006-1-AP), Mouse anti-Mitofusin (1:1000, Abcam, ab56889), Rabbit anti-TOM20 (1:2000, Cell Signaling Technology 42406), Rabbit anti-Histone H3 (1:1500, Abcam), Mouse anti-uL1 (1:1000 Santa Cruz, sc-100827). Secondary antibodies used were mouse anti-goat IgG–HRP (1:5000, Santa Cruz sc-2354), ECL Sheep anti-mouse IgG-HRP (1:3000 GE Healthcare NA9310V) and ECL Donkey anti-rabbit IgG-HRP (1:3000, GE Healthcare NA9340V). Immunoblots were developed using Clarity Western ECL substrate (BioRad).

### RNA-Seq Sample Preparation

For each clonal cell line (3 lines expressing GST-3F and 3 lines expressing NOCT Δ(2-15)-3F), two 10 cm dishes were plated with 3×10^6^ cells in DMEM + 10% FBS and incubated for three days at 37 °C and 5% CO_2_. Cells were harvested by dissociation with TrypLE (Thermo), washed twice with cold 1x PBS, pH 7.4 (Gibco), and rapidly frozen at −80 °C. Expression of GST-3F and NOCT Δ(2-15)-3F was verified by western blotting using α-FLAG as described above. Total RNA was isolated from frozen cell pellets using a Maxwell RSC SimplyRNA Cells kit (Promega) using a 2x concentration of DNAse I. RNA was eluted in 50 µL nuclease-free water and was analyzed using a Bioanalyzer Tapestation (Agilent), with RIN values between 9.6 and 10.0. Total RNA (5 µg) was combined with ERCC Spike-In Standards (5 µl of 1:50 diluted stock solution; Invitrogen) and submitted to the University of Minnesota Genomics Center for library generation and sequencing. RNA TruSeq strand-specific 50 bp dual indexed RNA libraries were prepared using RiboZero rRNA depletion (Illumina) and sequenced on a HiSeq 2500 in High-output mode. Library generation and sequencing was performed by the University of Minnesota Genomics Center.

### RNA-Seq Data Analysis

Raw RNA-seq data for each sample contained between 34,593,128 and 40,679,504 paired-end reads, with both read 1 and 2 of length 51 b. We used FastQC with MultiQC (Ewels et al. 2016) to examine the quality of the sequencing reads (71,72). The average quality per position of the sequencing reads for each sample sequencing read file ranged from 29.65 to 37.38. For both read 1 and read 2 of each sample, bases at positions 1-10 showed a noticeable non-random overrepresentation of specific bases, likely coming from adaptor sequences. We trimmed the first 10 bases for both read 1 and read 2 for all samples using Cutadapt (version 1.8.1) to remove a fixed number of bases from the 5′ end of read 1 and 2 separately in single-end mode (-u 10) (73). We next performed a quality trimming using Trimmomatic (version 0.33) in paired-end mode requiring a minimum quality score for leading and trailing sequences of 3, a minimum average quality score of 15 within a window of 4 bases, and a minimum read length of 26 (LEADING:3 TRAILING:3 SLIDINGWINDOW:4:15 MINLEN:26). The minimum read length requirement is derived from STAR alignment recommendations (74,75). Trimming survival rates of paired-end reads for all samples were > 99%. Raw and processed data is deposited at NCBI GEO (GSE142749). Code used for RNA-seq data analysis is available at: https://github.com/freddolino-lab/2019_nocturnin.

### Alignments

For mapping of the sequencing reads, we build a reference for STAR (version 2.5.2b) that contains the human reference genome GRCh38 (release 92, accessed 2018-06-18), NOCT Δ(2-15)-3F pFC3F plasmid sequence, GST-3F pFC3F plasmid sequence, and the ERCC spike-ins. The STAR index was built from the customized combined genome with the above sequences as a FASTA file, annotations as a GTF file, and an overhang length of 100. STAR alignment was performed using trimmed reads 1 and 2 in paired-end mode for each sample and written as BAM files sorted by alignment coordinates (76). For all samples, > 91% of reads mapped uniquely as reported by STAR.

### Quantification

We quantified the features from alignment results using HTSeq (version 0.6.1p1) (Anders, Pyl, and Huber 2015), with stranded set to reverse and requirement of minimum alignment quality score set to 10, supplied with customized annotations as used in alignment (77). The overlap resolution of feature quantification in HTSeq was set as --mode=union with --nonunique=none, that is, counting overlaps of a feature with any position of a read towards the set of identified feature(s) of the corresponding read, and adding ambiguous feature counts to none of the identified feature(s). For all samples, the percentages of uniquely aligned read pairs assigned to non-ambiguous features range from 61.74% to 66.35%, covering 26221 to 27756 of the 58489 features in the input annotation.

### Differential expression analysis

We performed differential expression analysis using DESeq2 (version 1.16.1) on the gene features quantified by HTSeq as described above (78). The 3 samples with NOCT Δ(2-15)-3F over-expression were treated as biological replicates in the NOCT group, and the 3 samples with GST-3F over-expression as biological replicates for the GST group, with the GST group being the reference level in the subsequent differential expression analysis. Quantified gene features were pre-filtered by requiring a total of counts from all samples to be greater than 10, leaving us with 23417 features. We used the single-step wrapper function DESeq for default for standard differential expression analysis, and then used a patched version function lfcShrink for log fold change shrinkage, specifying the original DESeq2 shrinkage estimator. The patch applied to log fold change shrinkage does not alter the algorithm but adds a column of output for standard errors of log fold change. The 95% confidence intervals of RNA-seq log fold changes as shown in **Figure 5d** were calculated as log fold change plus or minus 1.96 times the standard error.

23,414 genes survived that had no count outliers (characterized by a non-NA p-value from DESeq2) and 17058 genes passed the independent filtering based on the mean of normalized counts in all samples for each feature (characterized by a non-NA p-value when interpreting DESeq2 results) (79). Significance was determined by an FDR-corrected p-value threshold of 0.05 and log fold change cutoffs as specified in the text. Statistics for all gene features passing the pre-filtering are reported in **Supporting Information Table S1**. The shrunken log2 fold changes of the NOCT group relative to the GST group were used as the gene expression change levels in Figure 5. In the MA plot (**Figure 5B**) showing the overall expression levels and expression change levels, the average expression levels were quantified as the mean of log2 transcript per million (TPM) of six samples, including each 3 replicates in the NOCT and GST groups, and expression changes were quantified as shrunken log2 fold changes. To calculate TPM, we used the median fragment length for each sample and the number of genomic positions that appear in at least one exon of an annotated transcript for the length of each gene. In the volcano plot (**Figure 5C**), the significance values were quantified as negative log10 of DESeq2 adjusted p-values. The volcano plot and the MA plot were generated with R (version 3.6.1), with major contributor packages including ggplot2 (version 3.1.1) and biomaRt (version 2.42.0).

### Pathway enrichment analysis

We used iPAGE to identify potential NOCT-regulated pathways as the informative GO terms in the NOCT sample group compared to the control GST group (40,80,81). First, a custom iPAGE database was built by retrieving gene and GO term information from Ensembl using the R package biomaRt (accessed 2018-08-27) including an “index” file containing all genes with matching GO terms, a “names” file of GO term details such as corresponding annotations and domains, and an empty “homologies” file (82-84). The custom database did not contain information of the over-expression components or ERCC spike-ins. iPAGE, with built-in procedures to ensure robustness, used the raw (unshrunken) log2 fold changes for all gene features as the input, distributed the expression changes into 9 equally sized bins, and enforced the default requirement of independence among detected GO terms. In Figure S2, we identified 7 GO terms of interest among iPAGE detected GO terms upon discussion in text. For each GO term, we listed all genes that are associated with the GO term according to the custom iPAGE database described above. We plotted the sorted DESeq2 shrunken log2 fold changes as the y-axis values for bars representing each gene. Genes with filled black bars were significant with adjusted p-values from DESeq2 less than 0.05.

### Reverse Transcription and Quantitative PCR

The MIQE reporting information for RT-qPCR assays is reported in **Supporting Information File**, along with sequences and details of primers. Three replicate samples of each of the clonal cell lines, which over-expressed either NOCT Δ(2-15)-3F or GST-3F produced for RNA-seq, were grown in T75 flasks at 37 °C and 5% CO_2_. Cells were trypsinized with TrypLE (Thermo) and resuspended in 10 mL DMEM + 10% FBS (Thermo). Cells were collected by centrifugation at 1000 x g and stored at −80 °C as pellets until further use. For the doxycycline-inducible cell lines (HEK293 TetON NOCT Δ(2-15)-3F and HEK293T Flp-In T-REx NOCT(1-431)-3F) and control parent lines (WT HEK293 cells, ATCC) and WT HEK293T Flp-In T-REx cells, Thermo) were seeded and treated with either DMSO (-Dox) or 1 µg/mL doxycycline in DMSO (+ Dox, Sigma) for 72 hours. Cells were collected and stored as described above. RNA was harvested from each cell line using the Maxwell RSC SimplyRNA Cells Kit (Promega). Briefly, cell pellets were resuspended in 200 µL of Maxwell Lysis buffer and then mixed with 200 µL of homogenization buffer with 1-Thioglycerol. Lysates were added to the Maxwell RSC SimplyRNA Cells Kit cartridges along with a 2x concentration of Maxwell DNaseI solution per cartridge. The Maxwell RSC SimplyRNA Cells protocol was run without modification on a Maxwell RSC Instrument (Promega) and RNA was eluted in 40 µL of RNase free water and stored at −80 °C.

RNA was quantified using absorbance at 260 nm on a Nanodrop One UV spectrophotometer (Thermo Fisher). Reverse transcription reactions were carried out using 500 ng of total RNA from each sample and the GoScript Reverse Transcription Mix, Random Primers Kit (Promega, A2800) according to the manufacturer’s protocol. For each sample, control reactions without reverse transcriptase (RT) were generated to assess background from other sources of DNA. RT+ and RT-reverse transcription reactions were diluted 67-fold in RNase/DNase free water and RT-qPCR was performed on the CFX96 Real Time System (BioRad) using GoTaq qPCR Master Mix (Promega). Cycling parameters were as follows: 1) 95 °C for 2 min, 2) 95 °C for 15 s, 3) variable annealing temperatures for 1 min, 4) 72 °C for 1 min. Steps 2-4 were repeated for a total of 40 cycles. Control reactions without template and without RT were performed for all primer sets. For each transcript, PrimeTime Assays (IDT) were used in qPCR experiments. For the cell lines generated for RNA-Seq, qPCR was performed with all of the listed primers. For the Doxycycline-inducible cell lines, qPCR was performed for NOCT and TBP transcript levels. Annealing temperatures for each primer set were optimized to 90-110 % efficiency at 200 nM primer concentration. Cycle threshold (Ct) values were measured using the CFX Manager software.

To provide a consistent analysis of the RT-qPCR data, we applied a Bayesian hierarchical model using the STAN programming language via the brms R package (85-87). Our approach closely follows that used in Arvola et al., we used as initial observables the efficiency-weighted Cq values of the NOCT and reference (TBP) transcripts within each experimental replicate (calculated as in Eq. 3 of (88,89). We then fitted a linear model with Student-t distributed errors in which we considered each Cq value to arise due to a linear combination of terms from the target of interest (NOCT or TBP), target:genotype interaction, target:genotype:induction interaction, target:date interaction (with date corresponding to the batch of which the sample was a part), and a group-level (random) effect for each biological replicate. Cq values were centered on zero prior to model fitting. We used a normal(0,3) prior for all population-level (fixed) effects, and brms default priors for all other parts of the model. The model was fitted using 6 Monte Carlo chains run for 5000 iterations each, and convergence assessed using standard brms/STAN convergence checks and manual inspection of the posterior predictive distributions.

### Cellular ATP assays

The following cell lines were used in these measurements: WT HEK293 (ATCC), HEK293 TetON NOCT Δ(2-15)-3F, WT HEK293T Flp-In T-REx cells (Thermo), and HEK293T Flp-In T-REx NOCT (1-431)-3F. HEK293 cell lines were plated in 6-well plates with 300,000 cells per well and the HEK293T Flp IN lines were plated at 500,000 cells per well in DMEM + 10% Tet-free FBS (Sigma) and 1 x GlutaMAX (Sigma). One full 6-well tissue culture plate was seeded for each cell line used. Cells were allowed to adhere for 24 hours before the induction of over-expression. For each cell line, 3 wells were treated with 1 µg/mL Doxycycline (Dox +, Sigma) in DMSO or with DMSO (Dox -). Cells were incubated for 72 hours post-induction before harvesting. Each well of HEK293 cells were harvested by treatment with 200 µL TrypLE (Invitrogen). Detached cells were resuspended with 800 µL of DMEM + 10% FBS and cells. HEK293T cells were resuspended in 3 mL of culturing media by pipetting. HEK293T cell suspensions were centrifuged at 1000 x g for 10 mins to collect cells and pellets were resuspended in 1 mL of DMEM+ 10% FBS. Cells were counted by mixing a sample of the cell suspension with 0.4% Trypan blue (Invitrogen) in a 1:1 ratio before counting on the Countess II cell counter (Invitrogen). For each sample, 20,000 viable cells were added to a 96-well white flat-bottomed plate (Nunclon). Each well was brought up to 100 µL of total volume with DMEM + 10% FBS. 3 wells of DMEM + 10% FBS alone were added for blank measurements. 100 µL of reconstituted Cell Titer Glo reagent (Promega) was added to each well and the plate was incubated for 10 minutes at room temperature before luminescence was measured using a GloMAX Discover Plate reader luminometer (Promega). Three replicate experiments were performed with three biological replicates for each cell type and treatment within each experiment for nine total measurements per cell type and treatment.

The cellular ATP assays were analyzed using Bayesian hierarchical models, following the same general approach as that outlined above for the analysis of RT-qPCR experiments. In this case we took as primary data the 0-centered log2 luminescence measurements output by the GloMAX assay, and fitted a model with population-level effects for the cell line, condition (induced vs. uninduced), and condition:cell line interaction; and group-level effects for the assay date, biological replicate, and date-treatment-cell line combination. We assumed Student-t distributed errors, with a normal(0,5) prior on the population-level effects and brms default priors for all other parameters. The model was fitted and assessed as described above for the RT-qPCR data.

### Cellular Metabolite assays

The following cell lines were used in these measurements: WT HEK293 (ATCC), HEK293 TetON NOCT Δ(2-15)-3F, WT HEK293T Flp-In T-REx cells (Thermo), and HEK293T Flp-In T-REx NOCT (1-431)-3F. HEK293 cell lines were plated in 6-well plates with 300,000 cells per well and the HEK293T Flp IN lines were plated at 500,000 cells per well in DMEM + 10% Tet-free FBS (Invitrogen) and 1x GlutaMAX (Invitrogen). One full 6-well tissue culture plate was seeded for each cell line used. Cells were allowed to adhere for 24 hours before the induction of over-expression. For each cell line, 3 wells were treated with 1 µg/mL Doxycycline (Dox +, Sigma) in DMSO or with DMSO (Dox -). Cells were incubated for 72 hours post-induction before harvesting. Each well of HEK293 cells were harvested by treatment with 200 µL TrypLE (Invitrogen). Detached cells were resuspended with 800 µL of DMEM + 10% FBS and cells. HEK293T cells were resuspended in 3 mL of culturing media by pipetting. All cell samples were centrifuged at 1000 x g for 10 mins to collect cells and pellets were resuspended in approximately 250 µL of 1x PBS. Cells were counted by mixing a sample of the cell suspension with 0.4% Trypan blue (Invitrogen) in a 1:1 ratio before counting on the Countess II cell counter (Invitrogen). For each sample, 100,000 viable cells were added to a 96-well white flat-bottomed plate (Nunclon) 2x wells were plated for each sample, one for use with the NADP^+^/NADPH Glo Assay and one for use in the NADH/NAD^+^ Glo Assay (Promega) such that all 4 metabolites will be measured on the same plate for each set of samples. Relative amounts of each metabolite were measured according to the manufacturer’s instructions. Briefly, each well was brought up to 50 µL of total volume with 1x PBS. 4 wells of 1x PBS were added for blank measurements to each plate, one per metabolite. Fifty µL of 1% DTAB in 0.2 N NaOH was added to each well to lyse cells (base treatment). Fifty µL of the 100 µL cell suspension in base treatment was removed and added to an empty well. Twenty-five µL of 0.4 N HCl was added to these samples (acid treatment). Plates were incubated at 60 C for 15 mins in order to selectively destroy the NADP^+^/NAD^+^ in the base treatment and the NADPH/NADH in the acid treatment. The samples were brought to room temperature before neutralization of the base solutions with 50 µL of 0.25 M Tris base +0.2N HCl and the acid solutions with 25 µL of 0.5 M Tris base. All samples are in 100 µL of total volume at this point, with 50,000 cells per well. 50 µL of sample was removed from the wells measuring NADPH, NADH, and NADP^+^, while 90 µL of volume was removed from the wells measuring NAD^+^ so that samples can be diluted into the linear range for the assay. Wells used for NAD^+^ measurements were brought back up to 50 µL total volume by adding 40 µL of 1 x PBS. Reconstituted NADP^+^/NADPH Glo reagent was added to one set of acid and base treated wells and reconstituted NAD^+^/NADH Glo reagent was added to the other set of acid and base treated samples. Plates were incubated at room temperature for 1 hour before luminescence was measured using a GloMAX Discover Plate reader luminometer (Promega). Three replicate experiments were performed with three biological replicates for each cell type and treatment within each experiment.

The metabolite assays were analyzed using Bayesian hierarchical models, following the same general approach as that outlined above for the analysis of RT-qPCR experiments and ATP measurements. We took as primary data the 0-centered log2 luminescence measurements output by the GloMAX assay and fitted a model with population-level effects for the cell line, metabolite of interest, condition (induced vs. uninduced), and all possible interactions of those three features; and group-level effects for the biological replicates. We assumed Student-t distributed errors, with a normal(0,5) prior on the population-level effects and brms default priors for all other parameters. The model was fitted and assessed as described above for the RT-qPCR data.

## Acknowledgements

The authors thank the members of the Goldstrohm, Freddolino, Trievel, and Rorbach laboratories for helpful suggestions and advice during the course of these studies. The Genotype-Tissue Expression (GTEx) Project was supported by the Common Fund of the Office of the Director of the National Institutes of Health, and by NCI, NHGRI, NHLBI, NIDA, NIMH, and NINDS. The data used for the analyses described in this manuscript were obtained from the GTEx Portal on 03/01/19.

---

**Supporting Information Figure S1.**
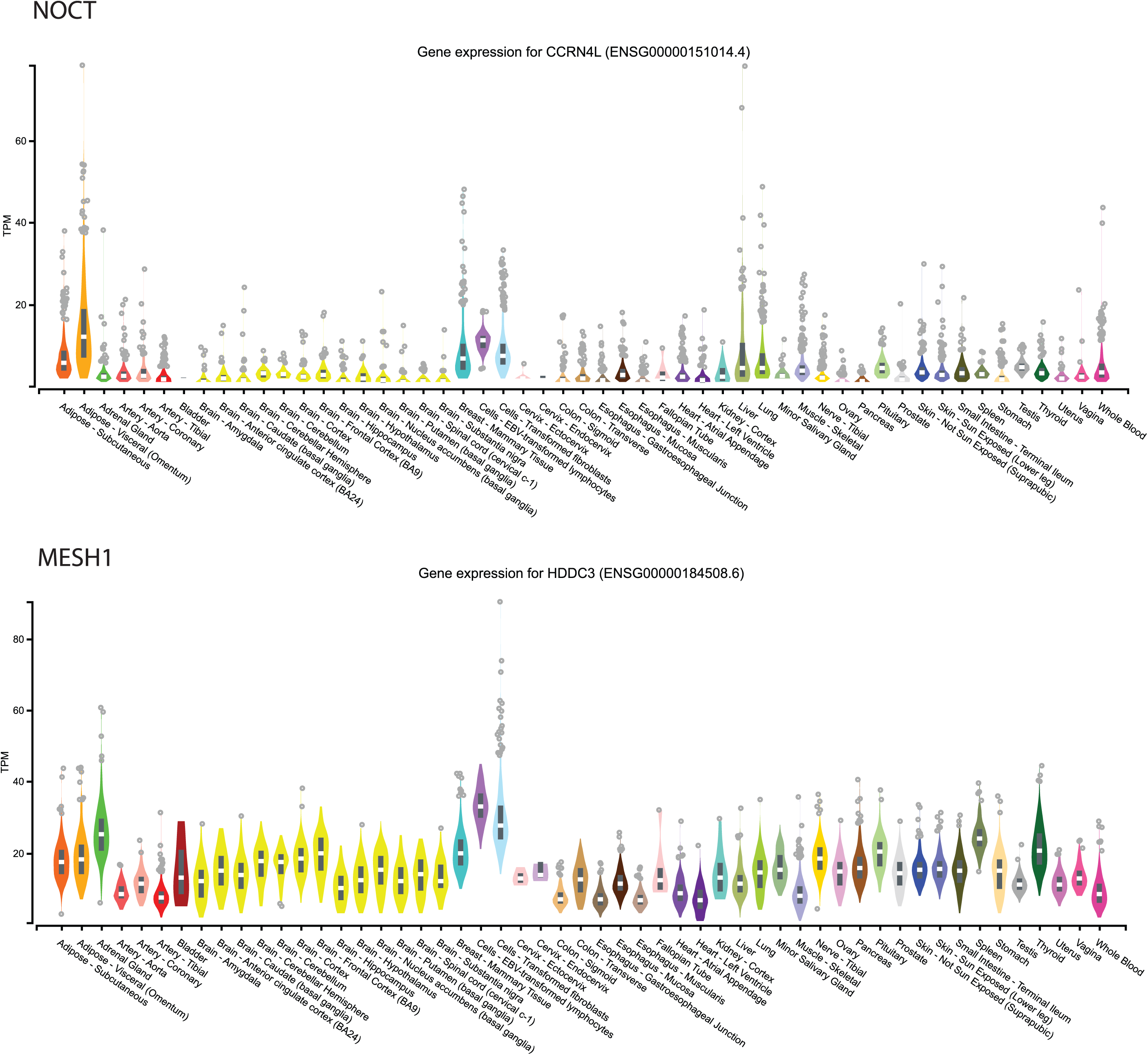
Expression of NOCT mRNA across different tissue types. Tissue specific mRNA expression data were obtained using the GTEx database with version 8 data with 17,382 RNA-Seq samples from 948 donors. Expression of *NOCT/CCRN4L* mRNA is plotted as transcripts per million (TPM) reads for each tissue type. *NOCT* expression is highest in adipose, liver, and mammary tissues. As a comparison, the expression of *MESH1/HDDC3* mRNA is also shown. *MESH1* is expressed more highly than *NOCT* in most tissue samples. As described by GTEx, values are calculated based on all isoforms from a single gene. The plots include mean expression level and distributions of the 25th and 75th percentiles with outliers shown as individual data points.

**Supporting Information Figure S2.**
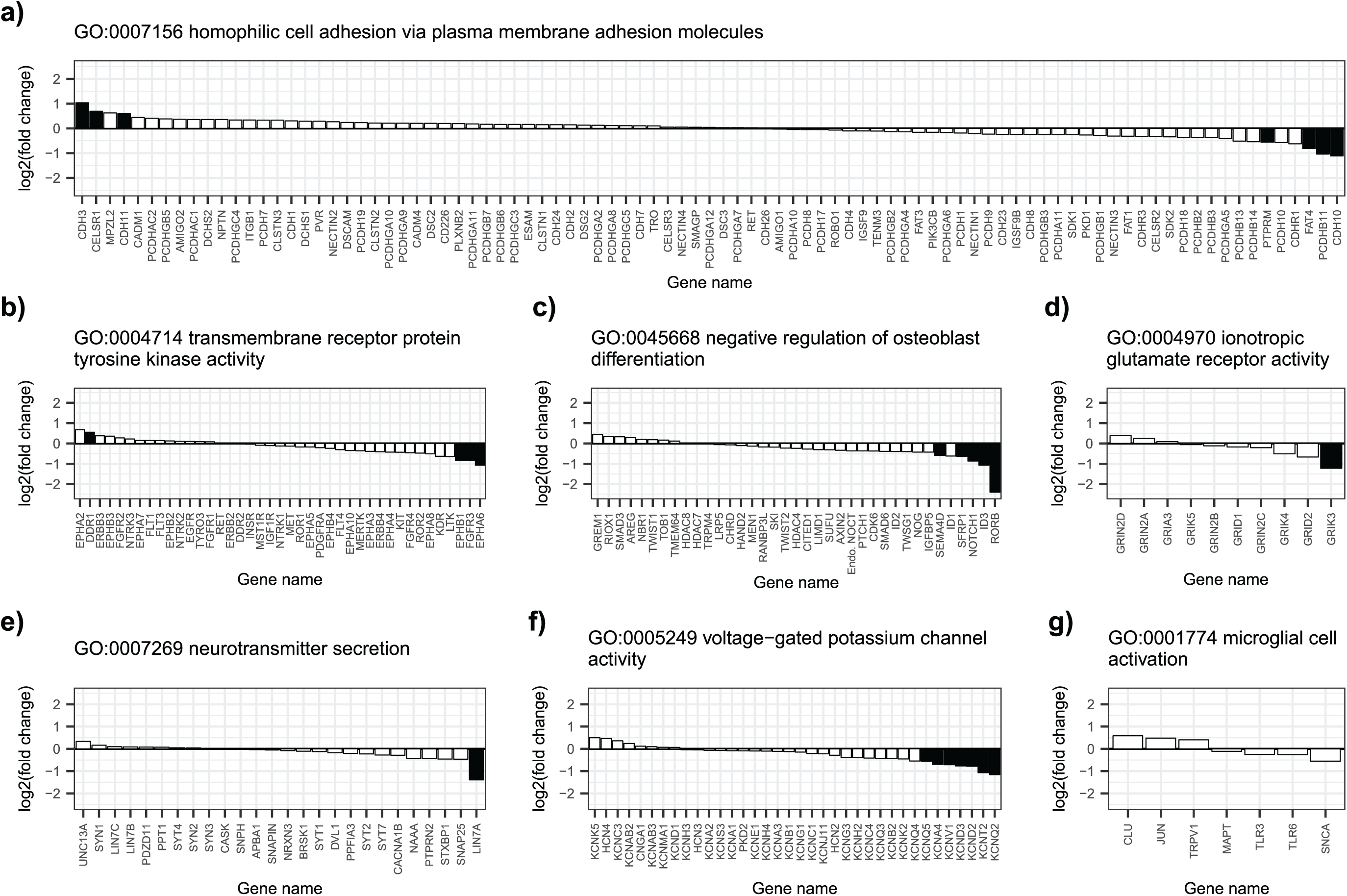
Gene level expression changes in gene ontology categories that are over-represented in NOCT Δ(2-15)-3F regulated RNA-seq dataset. For over-represented GO categories described in the main text, we identified the genes that are associated with the GO term and plotted the log2 fold changes of identified genes (in response to over-expressed NOCT Δ(2-15)-3F, relative to GST-3F) as the heights of bars in chart. The up-and down-regulation are reflected by the signs of log2 fold changes and subsequently the direction of bars in plots, in which the bars over zero as the y-axis show up-regulation of genes in response to NOCT Δ(2-15)-3F and bars below zero show down-regulation. Filled black bars indicate significance of gene expression changes by DESeq2. Data and statistics for each gene are reported in **Supporting Information Table S1**.

**Supporting Information Figure S3.**
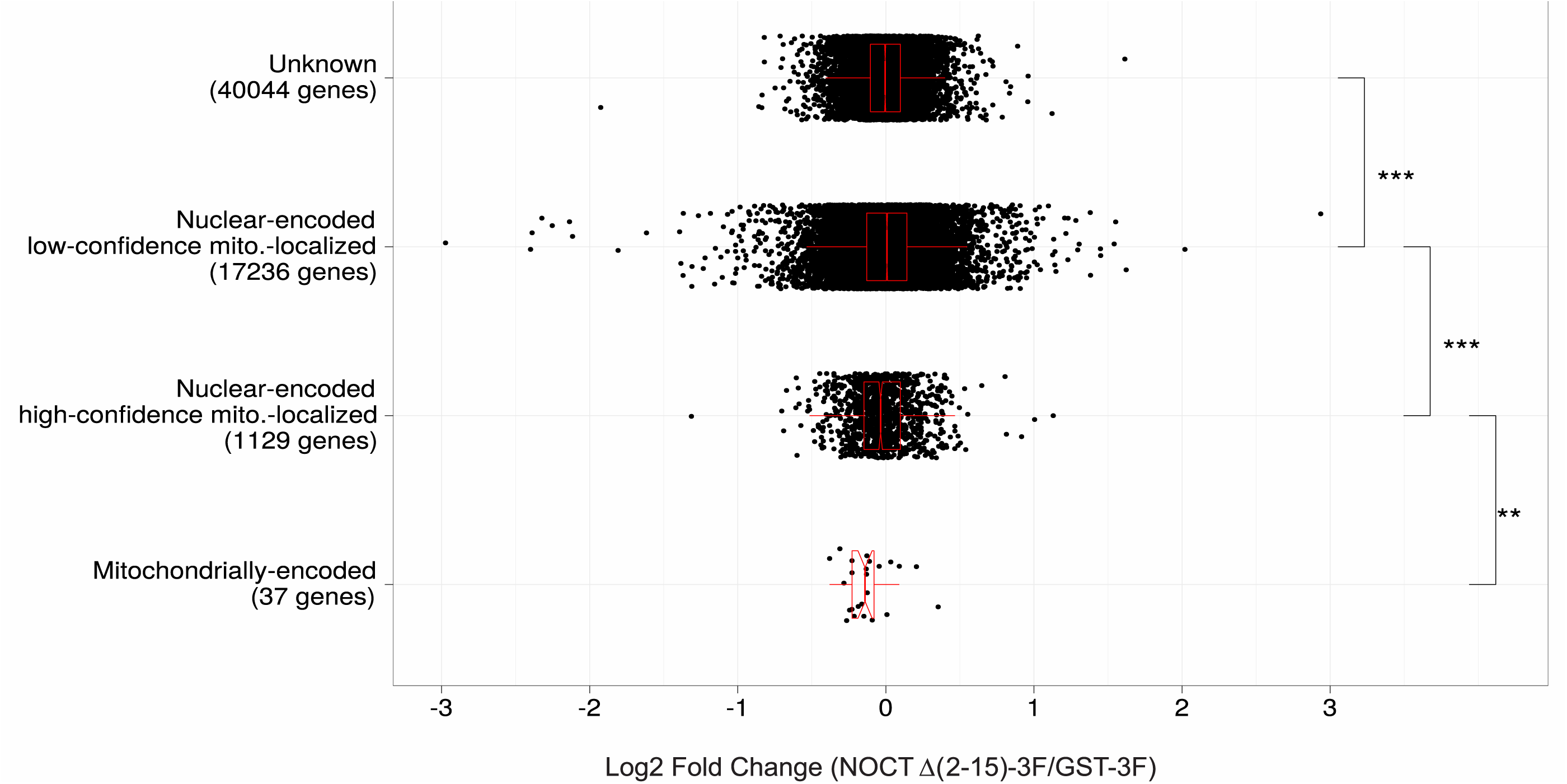
Distributions of NOCT Δ(2-15)-3F mediated changes of RNA levels from nuclear-or mitochondrially-encoded genes. Genes quantified in the RNA-seq experiment described in **Figure 5** are categorized as “mitochondrially-encoded genes”, “nuclear-encoded genes with high confidence of mitochondrial localization”, and “nuclear-encoded genes with low confidence of mitochondrial localization” based on the annotation from Human MitoCarta2.0; genes that are not found in MitoCarta2.0 are listed as “unknown” (90,91). Categorization of all genes is listed in **Supporting Information Table S1**. The number of genes in each category is noted in parentheses with the group label on y-axis. On the x-axis, the measure log2 fold change of genes in category in response to over-expressed NOCT Δ(2-15)-3F relative to negative control GST-3F is plotted. In the box plot, the lower and upper hinges correspond to the first and third quartiles, and the whiskers extend from the lower or upper hinges to the corresponding smallest or largest values no more than 1.5 times the inter-quartile range (IQR) more extreme from the hinge. The notches show the approximate 95% confidence intervals for group-level medians. Significances of group comparisons are determined by a two-sided Wilcoxon Rank Sum Test: *** = 0.001, ** = 0.01, * = 0.05.

**Supporting Information File. MIQE reporting data for Reverse Transcription and Quantitative Polymerase Chain Reaction experiments**

**Supporting Information Table S1. RNA seq data and statistics for transcriptome analysis in response to over-expressed NOCT Δ(2-15)-3F in HEK293 cells.**

**Supporting Information Table S2. Data and statistics for graphs in Figures 5 and 7.**

## References

1. Abshire, E. T., Chasseur, J., Bohn, J. A., Del Rizzo, P. A., Freddolino, P. L., Goldstrohm, A. C., and Trievel, R. C. (2018) The structure of human Nocturnin reveals a conserved ribonuclease domain that represses target transcript translation and abundance in cells. Nucleic acids research 46, 6257–6270

2. Baggs, J. E., and Green, C. B. (2003) Nocturnin, a deadenylase in Xenopus laevis retina: a mechanism for posttranscriptional control of circadian-related mRNA. Current biology : CB 13, 189–198

3. Blanco, A. M., Gomez-Boronat, M., Madera, D., Valenciano, A. I., Alonso-Gomez, A. L., and Delgado, M. J. (2017) First Evidences of Nocturnin in Fish: Two Isoforms in Goldfish Differentially Regulated by Feeding. Am J Physiol Regul Integr Comp Physiol, ajpregu 00241 02017

4. Green, C. B., Douris, N., Kojima, S., Strayer, C. A., Fogerty, J., Lourim, D., Keller, S. R., and Besharse, J. C. (2007) Loss of Nocturnin, a circadian deadenylase, confers resistance to hepatic steatosis and diet-induced obesity. Proceedings of the National Academy of Sciences of the United States of America 104, 9888–9893

5. Gronke, S., Bickmeyer, I., Wunderlich, R., Jackle, H., and Kuhnlein, R. P. (2009) Curled encodes the Drosophila homolog of the vertebrate circadian deadenylase Nocturnin. Genetics 183, 219–232

6. Hughes, K. L., Abshire, E. T., and Goldstrohm, A. C. (2018) Regulatory roles of vertebrate Nocturnin: insights and remaining mysteries. RNA Biol 15, 1255–1267

7. Goldstrohm, A. C., and Wickens, M. (2008) Multifunctional deadenylase complexes diversify mRNA control. Nature reviews. Molecular cell biology 9, 337–344

8. Douris, N., Kojima, S., Pan, X., Lerch-Gaggl, A. F., Duong, S. Q., Hussain, M. M., and Green, C. B. (2011) Nocturnin regulates circadian trafficking of dietary lipid in intestinal enterocytes. Current biology : CB 21, 1347–1355

9. Kawai, M., Green, C. B., Lecka-Czernik, B., Douris, N., Gilbert, M. R., Kojima, S., Ackert-Bicknell, C., Garg, N., Horowitz, M. C., Adamo, M. L., Clemmons, D. R., and Rosen, C. J. (2010) A circadian-regulated gene, Nocturnin, promotes adipogenesis by stimulating PPAR-gamma nuclear translocation. Proceedings of the National Academy of Sciences of the United States of America 107, 10508–10513

10. Kawai, M., Delany, A. M., Green, C. B., Adamo, M. L., and Rosen, C. J. (2010) Nocturnin suppresses igf1 expression in bone by targeting the 3’ untranslated region of igf1 mRNA. Endocrinology 151, 4861–4870

11. Garbarino-Pico, E., Niu, S., Rollag, M. D., Strayer, C. A., Besharse, J. C., and Green, C. B. (2007) Immediate early response of the circadian polyA ribonuclease nocturnin to two extracellular stimuli. Rna 13, 745–755

12. Estrella, M. A., Du, J., and Korennykh, A. (2018) Crystal Structure of Human Nocturnin Catalytic Domain. Scientific reports 8, 16294

13. Temme, C., Zhang, L., Kremmer, E., Ihling, C., Chartier, A., Sinz, A., Simonelig, M., and Wahle, E. (2010) Subunits of the Drosophila CCR4-NOT complex and their roles in mRNA deadenylation. Rna 16, 1356–1370

14. Wagner, E., Clement, S. L., and Lykke-Andersen, J. (2007) An unconventional human Ccr4-Caf1 deadenylase complex in nuclear cajal bodies. Molecular and cellular biology 27, 1686–1695

15. Kojima, S., Gendreau, K. L., Sher-Chen, E. L., Gao, P., and Green, C. B. (2015) Changes in poly(A) tail length dynamics from the loss of the circadian deadenylase Nocturnin. Scientific reports 5, 17059

16. Stubblefield, J. J., Gao, P., Kilaru, G., Mukadam, B., Terrien, J., and Green, C. B. (2018) Temporal Control of Metabolic Amplitude by Nocturnin. Cell reports 22, 1225–1235

17. Hee, S. W., Tsai, S. H., Chang, Y. C., Chang, C. J., Yu, I. S., Lee, P. C., Lee, W. J., Yun-Chia Chang, E., and Chuang, L. M. (2012) The role of nocturnin in early adipogenesis and modulation of systemic insulin resistance in human. Obesity 20, 1558–1565

18. Onder, Y., Laothamatas, I., Berto, S., Sewart, K., Kilaru, G., Bordieanu, B., Stubblefield, J. J., Konopka, G., Mishra, P., and Green, C. B. (2019) The Circadian Protein Nocturnin Regulates Metabolic Adaptation in Brown Adipose Tissue. iScience 19, 83–92

19. Pearce, S. F., Rorbach, J., Van Haute, L., D’Souza, A. R., Rebelo-Guiomar, P., Powell, C. A., Brierley, I., Firth, A. E., and Minczuk, M. (2017) Maturation of selected human mitochondrial tRNAs requires deadenylation. Elife 6

20. Rorbach, J., Nicholls, T. J., and Minczuk, M. (2011) PDE12 removes mitochondrial RNA poly(A) tails and controls translation in human mitochondria. Nucleic acids research 39, 7750–7763

21. Wood, E. R., Bledsoe, R., Chai, J., Daka, P., Deng, H., Ding, Y., Harris-Gurley, S., Kryn, L. H., Nartey, E., Nichols, J., Nolte, R. T., Prabhu, N., Rise, C., Sheahan, T., Shotwell, J. B., Smith, D., Tai, V., Taylor, J. D., Tomberlin, G., Wang, L., Wisely, B., You, S., Xia, B., and Dickson, H. (2015) The Role of Phosphodiesterase 12 (PDE12) as a Negative Regulator of the Innate Immune Response and the Discovery of Antiviral Inhibitors. The Journal of biological chemistry 290, 19681–19696

22. Estrella, M. A., Du, J., Chen, L., Rath, S., Prangley, E., Chitrakar, A., Aoki, T., Schedl, P., Rabinowitz, J., and Korennykh, A. (2019) The Metabolites NADP+ and NADPH are the Targets of the Circadian Protein Nocturnin (Curled). Biorxiv [Preprint]

23. Le, P. T., Bornstein, S. A., Motyl, K. J., Tian, L., Stubblefield, J. J., Hong, H. K., Takahashi, J. S., Green, C. B., Rosen, C. J., and Guntur, A. R. (2019) A novel mouse model overexpressing Nocturnin results in decreased fat mass in male mice. J Cell Physiol

24. Kozak, M. (1991) Effects of long 5’ leader sequences on initiation by eukaryotic ribosomes in vitro. Gene Expr 1, 117–125

25. Kozak, M. (1991) A short leader sequence impairs the fidelity of initiation by eukaryotic ribosomes. Gene Expr 1, 111–115

26. Aitken, C. E., and Lorsch, J. R. (2012) A mechanistic overview of translation initiation in eukaryotes. Nature structural & molecular biology 19, 568–576

27. Fukasawa, Y., Tsuji, J., Fu, S. C., Tomii, K., Horton, P., and Imai, K. (2015) MitoFates: improved prediction of mitochondrial targeting sequences and their cleavage sites. Mol Cell Proteomics 14, 1113–1126

28. Savojardo, C., Martelli, P. L., Fariselli, P., and Casadio, R. (2014) TPpred2: improving the prediction of mitochondrial targeting peptide cleavage sites by exploiting sequence motifs. Bioinformatics 30, 2973–2974

29. Indio, V., Martelli, P. L., Savojardo, C., Fariselli, P., and Casadio, R. (2013) The prediction of organelle-targeting peptides in eukaryotic proteins with Grammatical-Restrained Hidden Conditional Random Fields. Bioinformatics 29, 981–988

30. Bauer, N. C., Doetsch, P. W., and Corbett, A. H. (2015) Mechanisms Regulating Protein Localization. Traffic 16, 1039–1061

31. Shaw, G., Morse, S., Ararat, M., and Graham, F. L. (2002) Preferential transformation of human neuronal cells by human adenoviruses and the origin of HEK 293 cells. FASEB J 16, 869–871

32. Lin, Y. C., Boone, M., Meuris, L., Lemmens, I., Van Roy, N., Soete, A., Reumers, J., Moisse, M., Plaisance, S., Drmanac, R., Chen, J., Speleman, F., Lambrechts, D., Van de Peer, Y., Tavernier, J., and Callewaert, N. (2014) Genome dynamics of the human embryonic kidney 293 lineage in response to cell biology manipulations. Nature communications 5, 4767

33. Rasmussen, A. H., Rasmussen, H. B., and Silahtaroglu, A. (2017) The DLGAP family: neuronal expression, function and role in brain disorders. Mol Brain 10, 43

34. Byun, H., Lee, H. L., Liu, H., Forrest, D., Rudenko, A., and Kim, I. J. (2019) Rorbeta regulates selective axon-target innervation in the mammalian midbrain. Development 146

35. Roforth, M. M., Liu, G., Khosla, S., and Monroe, D. G. (2012) Examination of nuclear receptor expression in osteoblasts reveals Rorbeta as an important regulator of osteogenesis. J Bone Miner Res 27, 891–901

36. Mo, H. Y., Jo, Y. S., Yoo, N. J., Kim, M. S., Song, S. Y., and Lee, S. H. (2019) Frameshift mutation of candidate tumor suppressor genes QK1 and TMEFF2 in gastric and colorectal cancers. Cancer Biomark 24, 1–6

37. Xia, Z., Ouyang, D., Li, Q., Li, M., Zou, Q., Li, L., Yi, W., and Zhou, E. (2019) The Expression, Functions, Interactions and Prognostic Values of PTPRZ1: A Review and Bioinformatic Analysis. J Cancer 10, 1663–1674

38. Madeo, M., Stewart, M., Sun, Y., Sahir, N., Wiethoff, S., Chandrasekar, I., Yarrow, A., Rosenfeld, J. A., Yang, Y., Cordeiro, D., McCormick, E. M., Muraresku, C. C., Jepperson, T. N., McBeth, L. J., Seidahmed, M. Z., El Khashab, H. Y., Hamad, M., Azzedine, H., Clark, K., Corrochano, S., Wells, S., Elting, M. W., Weiss, M. M., Burn, S., Myers, A., Landsverk, M., Crotwell, P. L., Waisfisz, Q., Wolf, N. I., Nolan, P. M., Padilla-Lopez, S., Houlden, H., Lifton, R., Mane, S., Singh, B. B., Falk, M. J., Mercimek-Mahmutoglu, S., Bilguvar, K., Salih, M. A., Acevedo-Arozena, A., and Kruer, M. C. (2016) Loss-of-Function Mutations in FRRS1L Lead to an Epileptic-Dyskinetic Encephalopathy. Am J Hum Genet 98, 1249–1255

39. Schwenk, J., Boudkkazi, S., Kocylowski, M. K., Brechet, A., Zolles, G., Bus, T., Costa, K., Kollewe, A., Jordan, J., Bank, J., Bildl, W., Sprengel, R., Kulik, A., Roeper, J., Schulte, U., and Fakler, B. (2019) An ER Assembly Line of AMPA-Receptors Controls Excitatory Neurotransmission and Its Plasticity. Neuron 104, 680–692 e689

40. Goodarzi, H., Elemento, O., and Tavazoie, S. (2009) Revealing global regulatory perturbations across human cancers. Mol Cell 36, 900–911

41. Guntur, A. R., Kawai, M., Le, P., Bouxsein, M. L., Bornstein, S., Green, C. B., and Rosen, C. J. (2011) An essential role for the circadian-regulated gene nocturnin in osteogenesis: the importance of local timekeeping in skeletal homeostasis. Annals of the New York Academy of Sciences 1237, 58–63

42. Lehman, A., Thouta, S., Mancini, G. M. S., Naidu, S., van Slegtenhorst, M., McWalter, K., Person, R., Mwenifumbo, J., Salvarinova, R., Study, C., Study, E., Guella, I., McKenzie, M. B., Datta, A., Connolly, M. B., Kalkhoran, S. M., Poburko, D., Friedman, J. M., Farrer, M. J., Demos, M., Desai, S., and Claydon, T. (2017) Loss-of-Function and Gain-of-Function Mutations in KCNQ5 Cause Intellectual Disability or Epileptic Encephalopathy. Am J Hum Genet 101, 65–74

43. Niday, Z., Hawkins, V. E., Soh, H., Mulkey, D. K., and Tzingounis, A. V. (2017) Epilepsy-Associated KCNQ2 Channels Regulate Multiple Intrinsic Properties of Layer 2/3 Pyramidal Neurons. J Neurosci 37, 576–586

44. Kazak, L., Reyes, A., Duncan, A. L., Rorbach, J., Wood, S. R., Brea-Calvo, G., Gammage, P. A., Robinson, A. J., Minczuk, M., and Holt, I. J. (2013) Alternative translation initiation augments the human mitochondrial proteome. Nucleic acids research 41, 2354–2369

45. Kazak, L., Reyes, A., He, J., Wood, S. R., Brea-Calvo, G., Holen, T. T., and Holt, I. J. (2013) A cryptic targeting signal creates a mitochondrial FEN1 isoform with tailed R-Loop binding properties. PloS one 8, e62340

46. Lee, J., O’Neill, R. C., Park, M. W., Gravel, M., and Braun, P. E. (2006) Mitochondrial localization of CNP2 is regulated by phosphorylation of the N-terminal targeting signal by PKC: implications of a mitochondrial function for CNP2 in glial and non-glial cells. Mol Cell Neurosci 31, 446–462

47. Pearce, S. F., Rebelo-Guiomar, P., D’Souza, A. R., Powell, C. A., Van Haute, L., and Minczuk, M. (2017) Regulation of Mammalian Mitochondrial Gene Expression: Recent Advances. Trends Biochem Sci 42, 625–639

48. D’Souza, A. R., and Minczuk, M. (2018) Mitochondrial transcription and translation: overview. Essays Biochem 62, 309–320

49. Gilbert, M. R., Douris, N., Tongjai, S., and Green, C. B. (2011) Nocturnin expression is induced by fasting in the white adipose tissue of restricted fed mice. PloS one 6, e17051

50. Li, R., Yue, J., Zhang, Y., Zhou, L., Hao, W., Yuan, J., Qiang, B., Ding, J. M., Peng, X., and Cao, J. M. (2008) CLOCK/BMAL1 regulates human nocturnin transcription through binding to the E-box of nocturnin promoter. Mol Cell Biochem 317, 169–177

51. Kawai, M., Green, C. B., Horowitz, M., Ackert-Bicknell, C., Lecka-Czernik, B., and Rosen, C. J. (2010) Nocturnin: a circadian target of Pparg-induced adipogenesis. Annals of the New York Academy of Sciences 1192, 131–138

52. Ying, W. (2008) NAD+/NADH and NADP+/NADPH in cellular functions and cell death: regulation and biological consequences. Antioxid Redox Signal 10, 179–206

53. Xiao, W., Wang, R. S., Handy, D. E., and Loscalzo, J. (2018) NAD(H) and NADP(H) Redox Couples and Cellular Energy Metabolism. Antioxid Redox Signal 28, 251–272

54. Canto, C., Menzies, K. J., and Auwerx, J. (2015) NAD(+) Metabolism and the Control of Energy Homeostasis: A Balancing Act between Mitochondria and the Nucleus. Cell Metab 22, 31–53

55. Nikiforov, A., Dolle, C., Niere, M., and Ziegler, M. (2011) Pathways and subcellular compartmentation of NAD biosynthesis in human cells: from entry of extracellular precursors to mitochondrial NAD generation. The Journal of biological chemistry 286, 21767–21778

56. Adler, L. t., Chen, C., and Koutalos, Y. (2014) Mitochondria contribute to NADPH generation in mouse rod photoreceptors. The Journal of biological chemistry 289, 1519–1528

57. Davila, A., Liu, L., Chellappa, K., Redpath, P., Nakamaru-Ogiso, E., Paolella, L. M., Zhang, Z. G., Migaud, M. E., Rabinowitz, J. D., and Baur, J. A. (2018) Nicotinamide adenine dinucleotide is transported into mammalian mitochondria. Elife 7

58. Brooks, G. A. (2018) The Science and Translation of Lactate Shuttle Theory. Cell Metabolism 27, 757–785

59. Ohashi, K., Kawai, S., and Murata, K. (2012) Identification and characterization of a human mitochondrial NAD kinase. Nature communications 3, 1248

60. Kawai, S., and Murata, K. (2008) Structure and function of NAD kinase and NADP phosphatase: key enzymes that regulate the intracellular balance of NAD(H) and NADP(H). Biosci Biotechnol Biochem 72, 919–930

61. Ding, C. C., Rose, J., Wu, J., Sun, T., Chen, K., Chen, P., Xu, E., Tian, S., Akinwuntan, J., Guan, Z., Zhou, P., and Chi, J. A. (2018) Mammalian stringent-like response mediated by the cytosolic NADPH phosphatase MESH1. Biorxiv [Preprint]

62. Castello, A., Hentze, M. W., and Preiss, T. (2015) Metabolic Enzymes Enjoying New Partnerships as RNA-Binding Proteins. Trends in endocrinology and metabolism: TEM 26, 746–757

63. Hentze, M. W., Caughman, S. W., Rouault, T. A., Barriocanal, J. G., Dancis, A., Harford, J. B., and Klausner, R. D. (1987) Identification of the iron-responsive element for the translational regulation of human ferritin mRNA. Science 238, 1570–1573

64. Muckenthaler, M., Gray, N. K., and Hentze, M. W. (1998) IRP-1 binding to ferritin mRNA prevents the recruitment of the small ribosomal subunit by the cap-binding complex eIF4F. Mol Cell 2, 383–388

65. Mullner, E. W., Rothenberger, S., Muller, A. M., and Kuhn, L. C. (1992) In vivo and in vitro modulation of the mRNA-binding activity of iron-regulatory factor. Tissue distribution and effects of cell proliferation, iron levels and redox state. Eur J Biochem 208, 597–605

66. Constable, A., Quick, S., Gray, N. K., and Hentze, M. W. (1992) Modulation of the RNA-binding activity of a regulatory protein by iron in vitro: switching between enzymatic and genetic function? Proceedings of the National Academy of Sciences of the United States of America 89, 4554–4558

67. Dollenmaier, G., and Weitz, M. (2003) Interaction of glyceraldehyde-3-phosphate dehydrogenase with secondary and tertiary RNA structural elements of the hepatitis A virus 3’ translated and non-translated regions. J Gen Virol 84, 403–414

68. Nagy, E., Henics, T., Eckert, M., Miseta, A., Lightowlers, R. N., and Kellermayer, M. (2000) Identification of the NAD(+)-binding fold of glyceraldehyde-3-phosphate dehydrogenase as a novel RNA-binding domain. Biochem Biophys Res Commun 275, 253–260

69. Nagy, E., and Rigby, W. F. (1995) Glyceraldehyde-3-phosphate dehydrogenase selectively binds AU-rich RNA in the NAD(+)-binding region (Rossmann fold). The Journal of biological chemistry 270, 2755–2763

70. Minczuk, M., Piwowarski, J., Papworth, M. A., Awiszus, K., Schalinski, S., Dziembowski, A., Dmochowska, A., Bartnik, E., Tokatlidis, K., Stepien, P. P., and Borowski, P. (2002) Localisation of the human hSuv3p helicase in the mitochondrial matrix and its preferential unwinding of dsDNA. Nucleic acids research 30, 5074–5086

71. Andrews, S., Krueger, F., Segonds-Pichonn, A., Biggins, L., Krueger, C., and Wingett, S. (2010) FastQC: a quality control tool for high throughput sequence data.

72. Ewels, P., Magnusson, M., Lundin, S., and Kaller, M. (2016) MultiQC: summarize analysis results for multiple tools and samples in a single report. Bioinformatics 32, 3047–3048

73. Martin, M. (2011) Cutadapt Removes Adapter Sequences From High-Throughput Sequencing Reads. EMBnet.journal 17

74. Bolger, A. M., Lohse, M., and Usadel, B. (2014) Trimmomatic: a flexible trimmer for Illumina sequence data. Bioinformatics 30, 2114–2120

75. Dobin, A., Davis, C. A., Schlesinger, F., Drenkow, J., Zaleski, C., Jha, S., Batut, P., Chaisson, M., and Gingeras, T. R. (2013) STAR: ultrafast universal RNA-seq aligner. Bioinformatics 29, 15–21

76. Li, H., Handsaker, B., Wysoker, A., Fennell, T., Ruan, J., Homer, N., Marth, G., Abecasis, G., Durbin, R., and Genome Project Data Processing, S. (2009) The Sequence Alignment/Map format and SAMtools. Bioinformatics 25, 2078–2079

77. Anders, S., Pyl, P. T., and Huber, W. (2015) HTSeq--a Python framework to work with high-throughput sequencing data. Bioinformatics 31, 166–169

78. Love, M. I., Huber, W., and Anders, S. (2014) Moderated estimation of fold change and dispersion for RNA-seq data with DESeq2. Genome biology 15, 550

79. Bourgon, R., Gentleman, R., and Huber, W. (2010) Independent filtering increases detection power for high-throughput experiments. Proceedings of the National Academy of Sciences of the United States of America 107, 9546–9551

80. Ashburner, M., Ball, C. A., Blake, J. A., Botstein, D., Butler, H., Cherry, J. M., Davis, A. P., Dolinski, K., Dwight, S. S., Eppig, J. T., Harris, M. A., Hill, D. P., Issel-Tarver, L., Kasarskis, A., Lewis, S., Matese, J. C., Richardson, J. E., Ringwald, M., Rubin, G. M., and Sherlock, G. (2000) Gene ontology: tool for the unification of biology. The Gene Ontology Consortium. Nat Genet 25, 25–29

81. The Gene Ontology, C. (2019) The Gene Ontology Resource: 20 years and still GOing strong. Nucleic acids research 47, D330–D338

82. Zerbino, D. R., Achuthan, P., Akanni, W., Amode, M. R., Barrell, D., Bhai, J., Billis, K., Cummins, C., Gall, A., Giron, C. G., Gil, L., Gordon, L., Haggerty, L., Haskell, E., Hourlier, T., Izuogu, O. G., Janacek, S. H., Juettemann, T., To, J. K., Laird, M. R., Lavidas, I., Liu, Z., Loveland, J. E., Maurel, T., McLaren, W., Moore, B., Mudge, J., Murphy, D. N., Newman, V., Nuhn, M., Ogeh, D., Ong, C. K., Parker, A., Patricio, M., Riat, H. S., Schuilenburg, H., Sheppard, D., Sparrow, H., Taylor, K., Thormann, A., Vullo, A., Walts, B., Zadissa, A., Frankish, A., Hunt, S. E., Kostadima, M., Langridge, N., Martin, F. J., Muffato, M., Perry, E., Ruffier, M., Staines, D. M., Trevanion, S. J., Aken, B. L., Cunningham, F., Yates, A., and Flicek, P. (2018) Ensembl 2018. Nucleic acids research 46, D754–D761

83. Durinck, S., Spellman, P. T., Birney, E., and Huber, W. (2009) Mapping identifiers for the integration of genomic datasets with the R/Bioconductor package biomaRt. Nat Protoc 4, 1184–1191

84. Durinck, S., Moreau, Y., Kasprzyk, A., Davis, S., De Moor, B., Brazma, A., and Huber, W. (2005) BioMart and Bioconductor: a powerful link between biological databases and microarray data analysis. Bioinformatics 21, 3439–3440

85. Bürkner, P.-C. (2017) brms: An R Package for Bayesian Multilevel Models Using Stan. 2017 80, 28

86. Carpenter, B., Gelman, A., Hoffman, M. D., Lee, D., Goodrich, B., Betancourt, M., Brubaker, M., Guo, J., Li, P., and Riddell, A. (2017) Stan: A Probabilistic Programming Language. 2017 76, 32

87. Bürkner, P.-C. (2018) Advanced Bayesian Multilevel Modeling with the R Package brms. The R Journal 10, 395–411

88. Ganger, M. T., Dietz, G. D., and Ewing, S. J. (2017) A common base method for analysis of qPCR data and the application of simple blocking in qPCR experiments. BMC Bioinformatics 18, 534

89. Arvola, R. M., Chang, C. T., Buytendorp, J. P., Levdansky, Y., Valkov, E., Freddolino, P. L., and Goldstrohm, A. C. (2019) Unique repression domains of Pumilio utilize deadenylation and decapping factors to accelerate destruction of target mRNAs. Nucleic acids research

90. Calvo, S. E., Clauser, K. R., and Mootha, V. K. (2016) MitoCarta2.0: an updated inventory of mammalian mitochondrial proteins. Nucleic acids research 44, D1251–1257

91. Pagliarini, D. J., Calvo, S. E., Chang, B., Sheth, S. A., Vafai, S. B., Ong, S. E., Walford, G. A., Sugiana, C., Boneh, A., Chen, W. K., Hill, D. E., Vidal, M., Evans, J. G., Thorburn, D. R., Carr, S. A., and Mootha, V. K. (2008) A mitochondrial protein compendium elucidates complex I disease biology. Cell 134, 112–123

